# The successor representation in human reinforcement learning

**DOI:** 10.1101/083824

**Authors:** I Momennejad, EM Russek, JH Cheong, MM Botvinick, ND Daw, SJ Gershman

**Author notes:** Equal contribution (I Momennejad, EM Russek). Correspondence: Ida Momennejad. **Author contribution:** IM, EMR, MMB, NDD, and SJG designed the studies, JHC, EMR, and IM conducted the studies and collected data, IM, EMR, and SJG analyzed the data and ran model simulations, IM, EMR, MMB, NDD, and SJG wrote the manuscript.

## Abstract

Theories of reward learning in neuroscience have focused on two families of algorithms, thought to capture deliberative vs. habitual choice. “Model-based” algorithms compute the value of candidate actions from scratch, whereas “model-free” algorithms make choice more efficient but less flexible by storing pre-computed action values. We examine an intermediate algorithmic family, the successor representation (SR), which balances flexibility and efficiency by storing partially computed action values: predictions about future events. These pre-computation strategies differ in how they update their choices following changes in a task. SR’s reliance on stored predictions about future states predicts a unique signature of insensitivity to changes in the task’s sequence of events, but flexible adjustment following changes to rewards. We provide evidence for such differential sensitivity in two behavioral studies with humans. These results suggest that the SR is a computational substrate for semi-flexible choice in humans, introducing a subtler, more cognitive notion of habit.

## Introduction

How do humans and other animals discover adaptive behaviors in dynamic task environments? Identifying such behaviors poses a particular challenge in sequential decision tasks like chess or maze navigation, in which the consequences of an action may unfold gradually, over many subsequent steps and choices. In the past two decades, much attention has focused on a distinction between two families of reinforcement learning (RL) algorithms for solving multi-step problems, known respectively as *model-free* (MF) and *model-based* (MB) RL^1,2^. Although both approaches formalize the problem of choice as comparing the long-term future reward expected following different candidate actions, they differ in the representations and computations they use to estimate these values^3^ (Figure 1). The MF vs. MB dichotomy has been influential, in part, because it poses an appealingly clean tradeoff between decision speed and accuracy: MF algorithms store (“cache”) pre-computed long-run action values directly, whereas in MB algorithms, state transition learning enables more flexibility at greater computational expense: action values are recomputed using an internal model of one-step state transitions. This tradeoff has been put forward as a computational basis for phenomena relating to automaticity, deliberation, and control, and the inflexibility of MF learning in particular has been argued to explain maladaptive, compulsive behaviors such as drug abuse.

Although experiments suggest that people (and other animals) can flexibly alter their decisions in situations that would defeat fully MF choice, there remains surprisingly little evidence for how, or indeed whether, the brain carries out the sort of full MB re-computation that has typically been invoked to explain these capabilities. Furthermore, there exist other computational and representation learning shortcuts along the spectrum between MF and MB, which might suffice to explain many of the available experimental results. For the brain, such shortcuts provide plausible strategies for maximizing fitness; for the theorist, they enrich and complicate the theoretical tradeoffs involved in controlling decisions and managing habits. Here we report two experiments examining whether humans employ one important class of such shortcuts that lie between the MB and MF strategies. This intermediate algorithm for representation learning, based on the successor representation (SR)^4,5^, caches long-range or multi-step state predictions. In the remainder of the introduction we will focus on summarizing MF and MB algorithms and explaining SR-based algorithms in relation.

MF strategies, such as temporal difference (TD) learning, cache fully computed long-run *action values* as decision variables. Caching makes action evaluation at decision time computationally cheap, because stored action values can be simply retrieved. Action values (*Q* in Figure 1) can be estimated and updated using reward prediction error (RPE) signals and aggregating net value over a series of events and rewards unfolding over time. This means MF learners do not store any information about relations between different states, and hence fail to solve problems involving distal changes in reward value^6^. This has been empirically demonstrated by “reward revaluation” studies^7^ and latent learning^8^.

In contrast, MB algorithms do not rely on cached value functions. Instead, they store a full model of the world and compute trajectories at decision time. Specifically, they learn and store a one-step internal representation or model of the short-term environmental dynamics: specifically a state transition function T and a reward function R (T and R in Figure 1). By iterative computation using this one-step model, action values can be computed at decision time. This is analogous to mental simulation using a “cognitive map,” stringing together the series of outcomes expected to follow each action according to learned representations. This planning capacity endows MB algorithms with sensitivity to distal changes in reward (as in reward revaluation) and also to changes in the transition structure (such as detour problems in spatial tasks). This flexibility comes at a higher computational cost compared to caching: computations traversing a model are intensive in time and working memory. Such computation may be intractable (requiring error-prone approximations) in large search spaces such as wide and deep trees^9,10^.

The successor representation (SR) was originally introduced as a method for rapid generalization in reinforcement learning^4^. The SR simplifies evaluation via multi-step representation learning: it caches long-term predictions about the states it expects to visit in the future. Namely, for each starting state, the successor representation caches how often the agent expects or needs to visit each of its successor states in the future (which can be learned via simple temporal difference (TD) learning^5^). When the agent faces a decision, the successor representation is combined with the reward function (R) to evaluate optimal trajectory to reward: it combines how often successor states are expected to be visited on average in the future, with their reward.

Mathematically, the SR is a matrix *M*, where the *i*th row is a vector in which element, *M*(*i*, *j*), stores the expected discounted future occupancy of state *j* following initial state *i*. To understand what this means, imagine starting a trajectory in state *i*, and counting the number of times each state *j* is encountered subsequently, while exponentially discounting visits that occur farther in the future. This representation is useful because at decision time, action values can be computed by linearly combining the SR for the current state with the one-step reward function. This obviates the MB strategy’s laborious iterative simulation of future state trajectories using MB’s one-step model, but stops short of storing the fully computed decision variable, as does MF learning. Thus, action evaluation with the SR has similar computational complexity to MF algorithms, while at the same time retaining some of the flexibility characteristic of the MB strategy. This form of predictive caching, if it exists in the brain, would provide a compromise between fully flexible deliberation and complete automaticity, allowing choices to be adjusted nimbly in some circumstances but still producing inappropriate, habit-like behavior in others. Such representation learning strategy is particularly well-suited to environments where the trajectories of states are fairly reliable, but rewards and goals change frequently. In such “multi-goal” environments, as they are referred to by the RL literature, a compromise between MF and MB strategies becomes an appealing algorithm. The evidence that people and animals can solve (at least small and simple) reward revaluation tasks in spite of MB algorithms’ computational complexity lends further support to the validity of a more cost-efficient algorithm at play.

**Figure 1.**
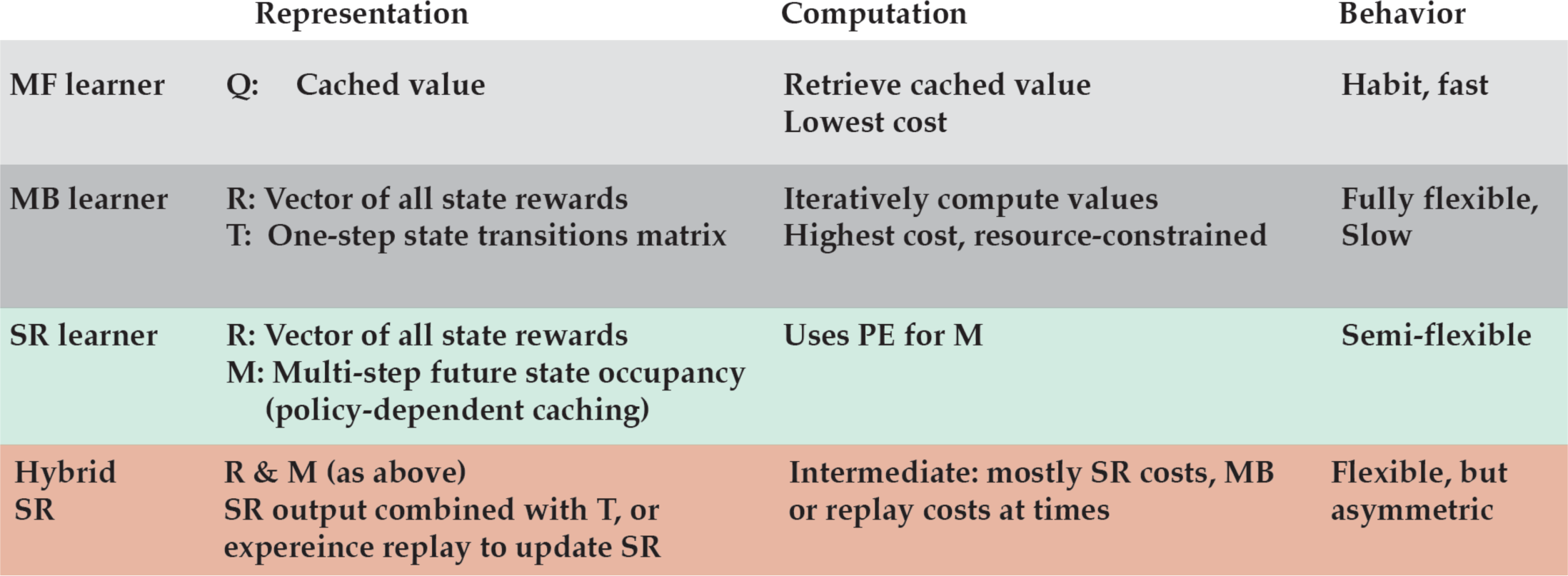
Comparison of stored representation, computations at decision time, and behavior across models. A brief comparison of the representations stored and used by different learning algorithms, their computational requirements at decision time, and their behavior. Q: value function (cached action values), R: reward function, T: full single-step transition matrix, M: the successor representation or a ‘rough’ predictive map of each state’s successor states. Both model-free cached value and the successor representation can be learned via simple temporal difference (TD) learning during the direct experience of trajectories in the environment.

**Figure 2.**
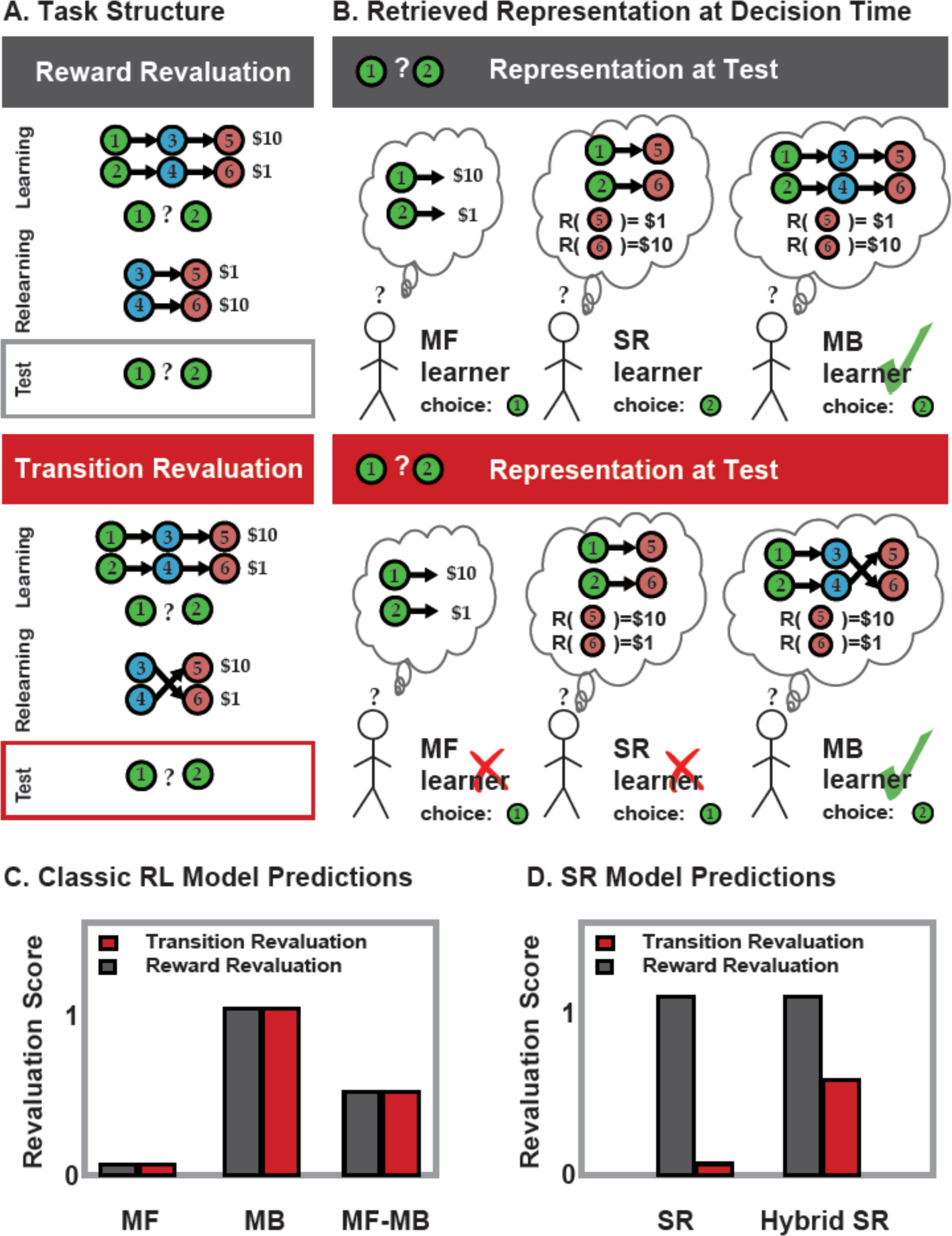
Schematic of retrieved representations at test and model predictions in reward and transition revaluation trials. (A) Schematics for task structures in reward revaluation (top) and transition revaluation (bottom). (B) Schematics of the representations retrieved at decision time by different learners: an MB learner retrieves 1-step transitions, and rolls out and computes full transitions (a costly computation), then combines them with the reward vector R to produce a decision. The MB learner is equally successful in both reward and transition revaluation. A purely SR learner retrieves the successor representation (of which only the relevant rows are displayed here) without further computation and readily combines it with the reward vector. (C) Qualitative model predictions for reward (gray) and transition (red) revaluation. Predicted revaluation scores for model-free (MF), model based (MB), mixture model (MB-MF), purely successor representation (SR), and a hybrid SR learner (which would combine SR with model-based or replay, hybrid SR-MB and SR-replay for short). Classic RL solutions all predict symmetrical responses to the retrospective revaluation problems. That is, while the model-free strategy has no solution to reward or transition revaluations, MB and hybrid MF-MB learners predict symmetrical performance for all types of revaluation. (D) successor representation (SR) strategies predict asymmetrical responses: the SR algorithm is sensitive to changes in reward. However, since SR stores a multi-step predictive map M, and not the step-by-step transition structure, it cannot update M in absence of direct experience. That is, the SR effectively “compiles” the transition structure into an aggregate predictive representation of future states, and therefore, cannot adapt to local changes in the transition structure in the environment, without experiencing the new trajectories in full. While a pure SR algorithm cannot solve transition revaluation, a hybrid SR learner that is updated via simulated experience, e.g. via MB representations or episodic replay, adjusts its decision for any revaluation, but performs best in reward revaluation.

Several lines of evidence motivate consideration of the SR as a hypothesis for biological reinforcement learning. First, converging evidence from other domains suggests that the SR is explicitly represented in the brain: the SR defined over space can capture many properties of rodent hippocampal place cells^11^, whereas in tasks with more abstract sequential stimulus structure, the SR captures properties of fMRI pattern similarity in hippocampal and prefrontal areas^12,13^. The SR is also closely related to the representation posited by the Temporal Context Model of memory^5^, which successfully explains numerous memory effects. Second, SR learning can be implemented using a version of the predominant neural theory of MF learning, the TD learning theory of the dopamine response^14^. In particular, if the expected future visits stored in M are used as the state input for a TD learner, TD will learn the reward function R^4^. Together with the MB-like flexibility of the SR, this observation may help to explain several puzzling reports that dopamine affects putative behavioral signatures of not just MF but also MB learning^15,16^. These results are unexpected for standard MB algorithms, which do not share any aspects of their learning with TD. Third, most existing empirical evidence in favor of MB algorithms in the brain is equally consistent with an SR-based account. In particular, SR and MB algorithms make identical predictions about reward revaluation experiments, a class which includes reward devaluation, latent learning^8,17^, sensory preconditioning^18,19^, and the two-step Markov decision tasks that have been widely used with humans^6,15^. Discriminating between these accounts, however, stands as an important empirical challenge that is taken up by the present paper.

In the present work, we provide new experimental designs that aim to tease apart behavior using SR and MB computations, and in particular to investigate whether people learn and use representations that cache long run expectancies about future state occupancy. Although the SR can flexibly adapt to distal changes in reward (as in reward revaluation), it cannot do so with distal changes in the transition structure (what we call transition revaluation). Because the SR caches a predictive representation that effectively aggregates over the transition structure, it cannot be flexibly updated in response to changes in this structure, unlike an MB strategy. Instead, the SR can only learn about changes in the transition structure incrementally and through direct experience, much like the way MF algorithms learn about changes in the reward structure. We exploited this difference by comparing the effects of reward and transition revaluation manipulations on human behavior. MF algorithms predict that participants will be equally insensitive to reward and transition revaluation, whereas MB algorithms predict that participants will be equally sensitive to both (and accordingly any linear combination of the two algorithms will predict equal sensitivity to both conditions). Crucially, any learning strategy that uses the SR (either SR alone or a hybrid SR strategy that combines SR with the other strategies) predicts that participants will be more sensitive to reward than transition devaluation (Figure 2).

To summarize, MF strategies do not store any representations of future states and do not compute state representations at decision time (Figures 1 and 2). MB strategies, on the other hand, store and retrieve one-step representations, leading to high computational demand at decision time. However, the SR caches a predictive map of states that the agent expects to visit in the future. Using these cached representations at decision time, the SR can solve reward revaluation but not transition revaluation, while MB is equally successful at all revaluations and MF is equally unsuccessful. Another possibility is to have a blend of SR with other strategies, which we will refer to as hybrid SR strategies. Hybrid SR strategies could combine the predictive representation with MB computations or replay in order to either update the SR offline in a Dyna-like architecture^20^ (which we call SR-Dyna) or augment the SR at decision time (which we refer to as hybrid SR-MB or SR-replay strategies). As such, all hybrid SR strategies will perform better than a pure SR strategy on transition revaluation (but worse than MB). Specifically, hybrid SR strategies predict higher accuracy and faster reaction times for reward revaluation than transition revaluation (an asymmetry in performance that is not predicted by either MF or MB; see Figures 1 and 2). We experimentally test and confirm these predictions in two studies, providing direct evidence for the SR in human reinforcement learning.

## Results

### Experiment 1: Differential sensitivity to reward and transition revaluation in a passive learning task

We designed a multistep sequential learning task to compare human behavior under reward revaluation and transition revaluation. Experiment 1 used a passive learning task, which permitted the simplest possible test of the theory, removing the need to model action selection. A schematic of the design is displayed in Figure 3 and Supplementary Figure 1. Participants played twenty games, each of which was made up of three phases. In Phase 1 (the learning phase), participants first learned three-step trajectories leading to reward. These trajectories were deterministic and passively experienced (i.e., transition required no action from the participant). Participants were exposed to one stimulus at a time, and were asked to indicate their preference for the middle state after every five stimuli. The learning phase ended if the participant indicated preference for the highest paying trajectory three times, or after 20 stimulus presentations. At the end of the learning phase, they were asked to indicate which starting state they believed led to greater future reward by reporting their relative preference using a continuous scale. Learning was assessed by the participant’s preference for the starting state associated with the more rewarding trajectory.

**Figure 3.**
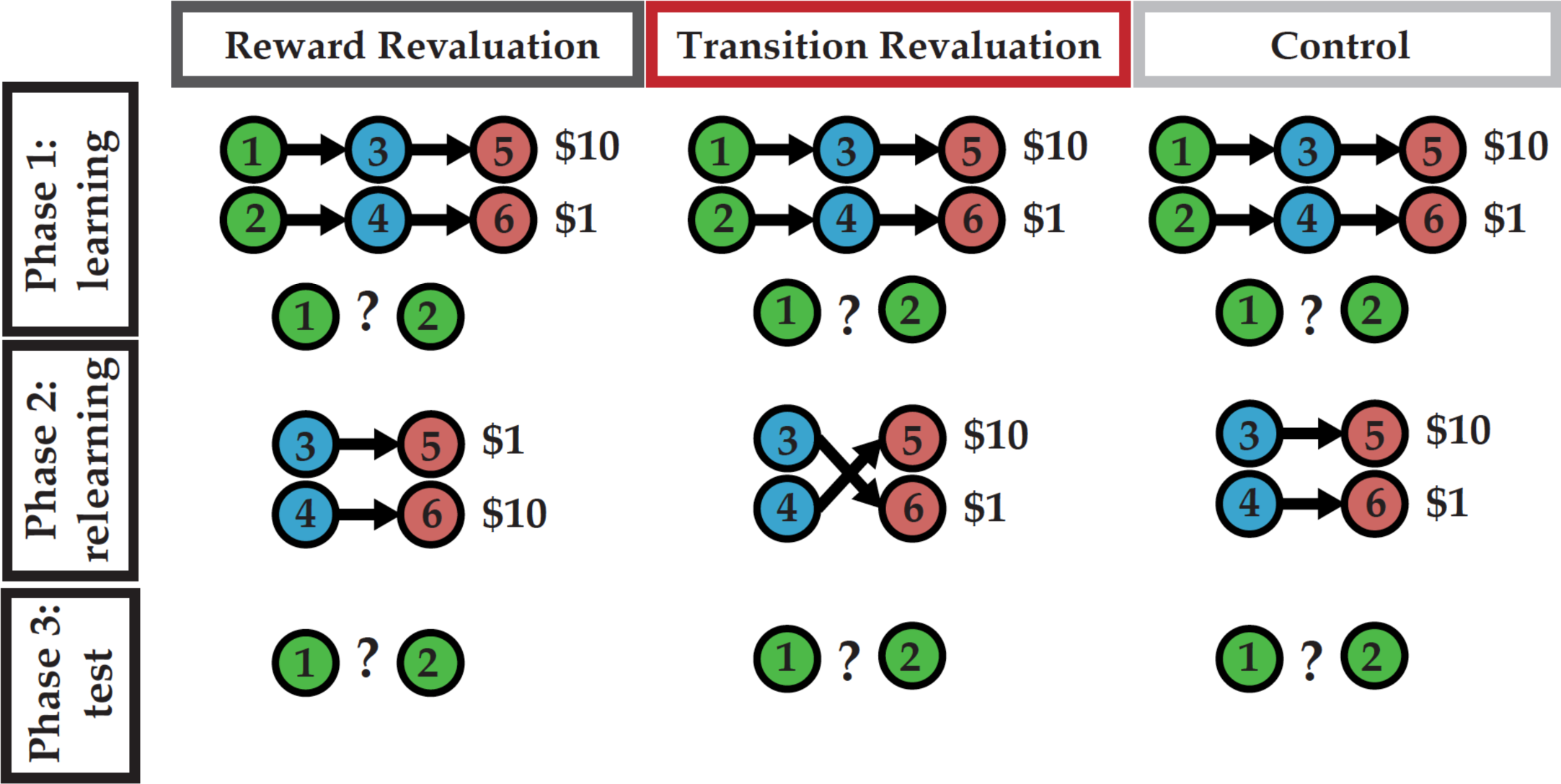
Schematic of Study I experimental design. A schematic of experimental conditions with reward and transition revaluation and a control condition. The graph represents the structure of states, rewards, and revaluation conditions. Participants never saw these graphs, and experienced the task structure one stimulus at a time as displayed in B. The experiment consisted of three phases: learning, relearning/revaluation, and test in extinction. At the end of phase 1 (learning) and phase 3 (test), participants indicated their preference for the starting state of the trajectory that would maximize reward. Revaluation scores were calculated as the difference between participants’ preference scores phases 1 and 3.

In Phase 2 (the relearning phase), trajectories were initiated at the second or middle state of the trajectory, and the structure of the task was altered in one of two ways (manipulated within participant, across games): in the reward revaluation condition, the rewards associated with the terminal states were swapped, whereas in the transition revaluation condition, the transitions between step 2 and step 3 states were swapped (Figures 2 and 3). Both conditions induced equivalent changes to the values of the first-stage states. In addition, we included a control condition in which no change occurred during phase 2 (the relearning phase). As in phase 1, participants were probed for state 2 preferences after each 5 stimuli, and phase 2 ended if participants had indicated the middle state of the most rewarding trajectory 3 times or after 20 stimuli (see Methods and Supplementary Figure 1). Finally, in phase 3 (test phase), participants were again asked which starting state they preferred. They had 20 s to give a response. Revaluation was measured as the amount of preference change between phases 1 and 3 (Δpreference, signed so that positive values indicate preference shift toward the newly optimal starting state; Figure 3, Supplementary Figure 1). See *Methods* for a more detailed explanation of the experimental design.

Figure 4A displays mean (+/− 1 SEM) revaluation scores for the three conditions, *n = 58 participants* (Reward Revaluation: *mean = .5199, SEM = .0203*, Transition Revaluation: *mean = .4503, SEM = .0229*, Control: *mean = .0310, SEM = .0310*). Revaluation scores were significantly higher for the reward revaluation than for both the transition revaluation [*t(57) = 2.89, p < 0.01*] and the control condition [*t(57) = 10.14, p < 0.001*]. Furthermore, revaluation scores were significantly higher in the transition revaluation compared to the control condition [*t(57) = 9.05, p < 0.001*]. In the control condition, as expected, revaluation scores were not significantly different from zero, [*t(57)=1.2; p = 0.22*]. This finding is important because it verifies that baseline forgetting or randomness cannot explain participants’ behavior in the non-control conditions. We conducted a repeated-measures ANOVA and post hoc t-tests (on participant means) using Bonferroni correction. A one-way within-participants ANOVA revealed a significant effect of condition on revaluation scores: F(2, 1154) = 38.77, P < .001. Because our experiment was designed with clear *a priori* hypotheses, we performed planned comparisons using paired sample t-tests with Bonferroni correction, which indicated that mean revaluation score in reward revaluation condition was significantly different than transition revaluation condition: t(57) = 2.8928, p = 0.016. In addition, mean revaluation scores were higher in both reward revaluation (t(57) = 10.148, p < .001) and transition revaluation (t(57) = 9.0543, p < .001) than the control condition.

We further analyzed the data for any time-on-task effects on accuracy or differences in accuracy, i.e. whether behavior improved as a result of practice, and whether these changes were significantly different in transition vs. reward revaluation conditions. For non-control trials, there was a significant effect of time on task (trial number) on the revaluation score [*F(1, 57.259) = 9.9171, P < 0.01*] indicating that participants improved at the task over time. However, there was no significant interaction of this effect with revaluation condition [*F(1, 68.284) = .15436, P = 0.695*]. Finally, we also found a significant main effect of revaluation condition on response times during the test phase [*F(2, 171) = 7.74, p < 0.001*; Figure 4A, right]. In particular, response times were slower in the transition revaluation condition compared to both the reward revaluation condition [*t(57) = 2.08, p < 0.05*] and the control condition [*t(57) = 4.04, p < 0.001*], and response times in the reward revaluation condition were significantly slower compared to the control condition where no changes had occurred [*t(57) = 3.5646, p < 0.001*]. There were substantial individual differences in the behavior under different revaluation conditions, and the appearance of multimodality (Figure 4B). Previous research has suggested that the balance between model-based vs. model-free learning tracks important individual differences, such as symptoms of mental illness ^21^. Future work exploring individual differences in subtler forms of computation and representation, presented here, may provide valuable insights into the arbitration of different representation and control processes across populations, as well as their effect on the flexibility and pathologies of decision-making.

**Figure 4.**
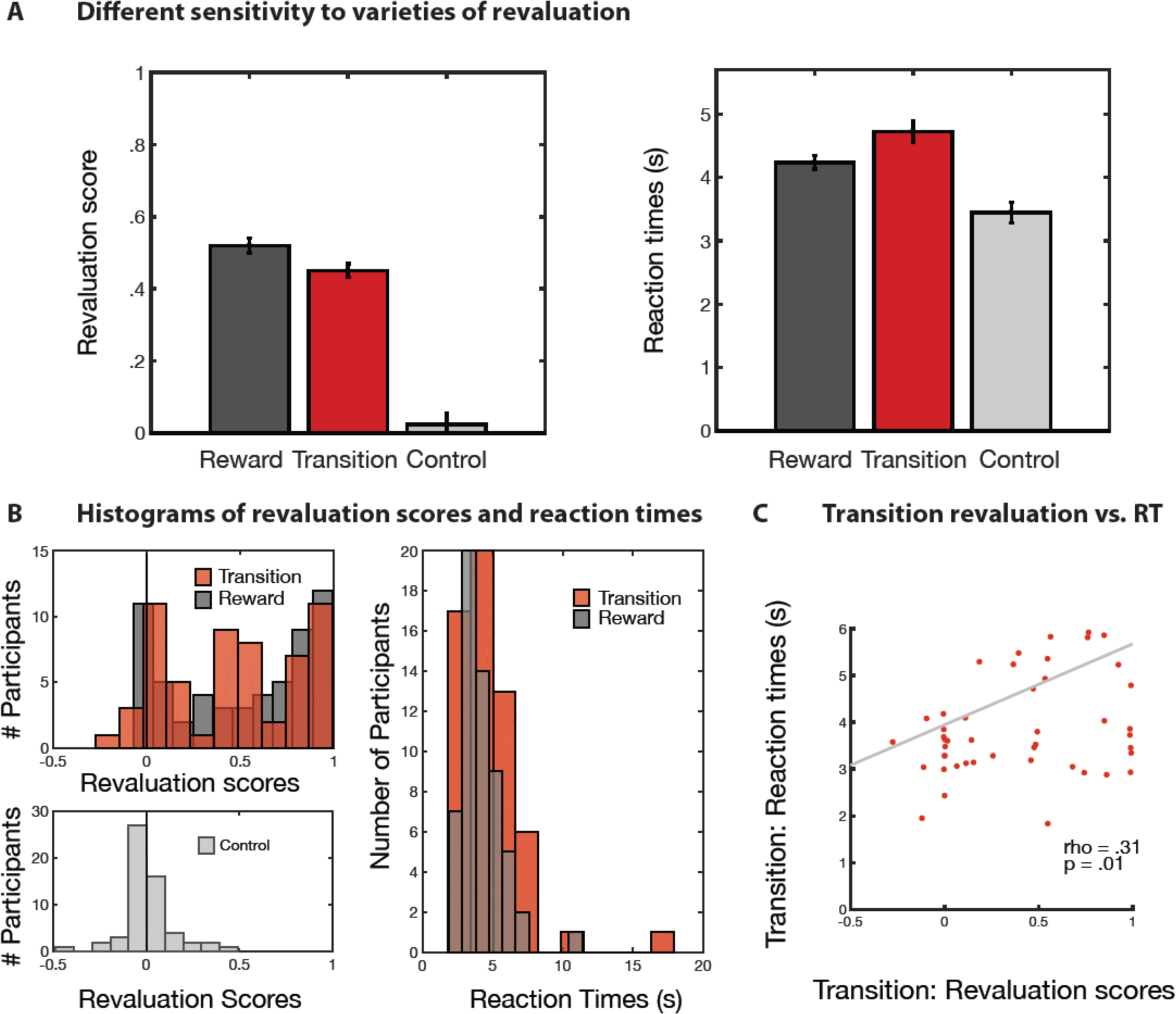
Behavioral performance in a passive sequential learning task. Human performance was measured as the change in their preference ratings for the starting states. Revaluation scores denote the change in a given game’s relative preference rating, after vs. before the relearning phase. (A, left) Mean revaluation scores are plotted for the three conditions: reward revaluation, transition revaluation, and control games. There was a significant main effect of condition. (A, right) Mean reaction times (s) to the final preference decision in phase 3 under reward and transition revaluation. Behavioral responses to the preference rating were significantly slower during transition revaluation. Error bars indicate SEM. (B) Histograms reveal the distribution of revaluation scores and response times across the main conditions. (C) There was a significant correlation between the accuracy of transition revaluation responses and reaction times: more accurate transition revaluation took longer, suggesting that successful transition revaluation might have involved more computation at decision time. Together with significantly higher reaction times compared to reward revaluation (A, right), this positive correlation lends further evidence to the possibility that compared to reward revaluation, transition revaluation required more cycles of computation at decision time, relying less on cached representations.

### A hybrid model that combines model-based learning with the successor representation explains differential sensitivity to varieties of revaluation

The key signature of SR’s caching of multistep future state occupancies (i.e. caching how often the agent expects or needs to visit a successor state in the future) is differential sensitivity to reward vs. transition revaluations. Participants’ differential sensitivity to these manipulations argues against a pure MB or MF account (see *Methods* for a detailed description of all the models considered here). MF algorithms predict equivalent and total insensitivity to both revaluation conditions, because participants are never given the opportunity to re-experience the start state following the revaluation phase. This effectively fools algorithms like TD learning that rely on chaining of trajectories of direct experience to incrementally update cached value estimates. In contrast, MB algorithms predict equal sensitivity to both conditions (Figure 5, left panel) so long as the revalued contingencies are themselves learned, because the updated internal model following the revaluation phase will produce accurate action values for the start state in either case. Accordingly, any weighted combination of these two evaluation mechanisms – which is the hybrid reinforcement learning model often used to explain previous sequential decision tasks^22^ – also does not predict differential sensitivity. This is because the combination will simply scale the equal sensitivity of either algorithm up or down.

SR-based algorithms fare better (Figure 5, middle panel), in that they predict that an agent will be insensitive to transition revaluation but sensitive to reward revaluation. In particular, algorithms that update a cached estimate of the SR using a TD-like learning rule require full trajectories through the state-space in order to update the start state’s SR (i.e., the future state occupancies it predicts) following the revaluation phase. This mirrors the direct experience requirement of MF algorithms for value estimation. However, unlike MF algorithms, SR-based algorithms can instantly adapt to changes in reward structure, because this only requires updating the immediate reward prediction, which then propagates through the entire state space when combined with the SR.

**Figure 5.**
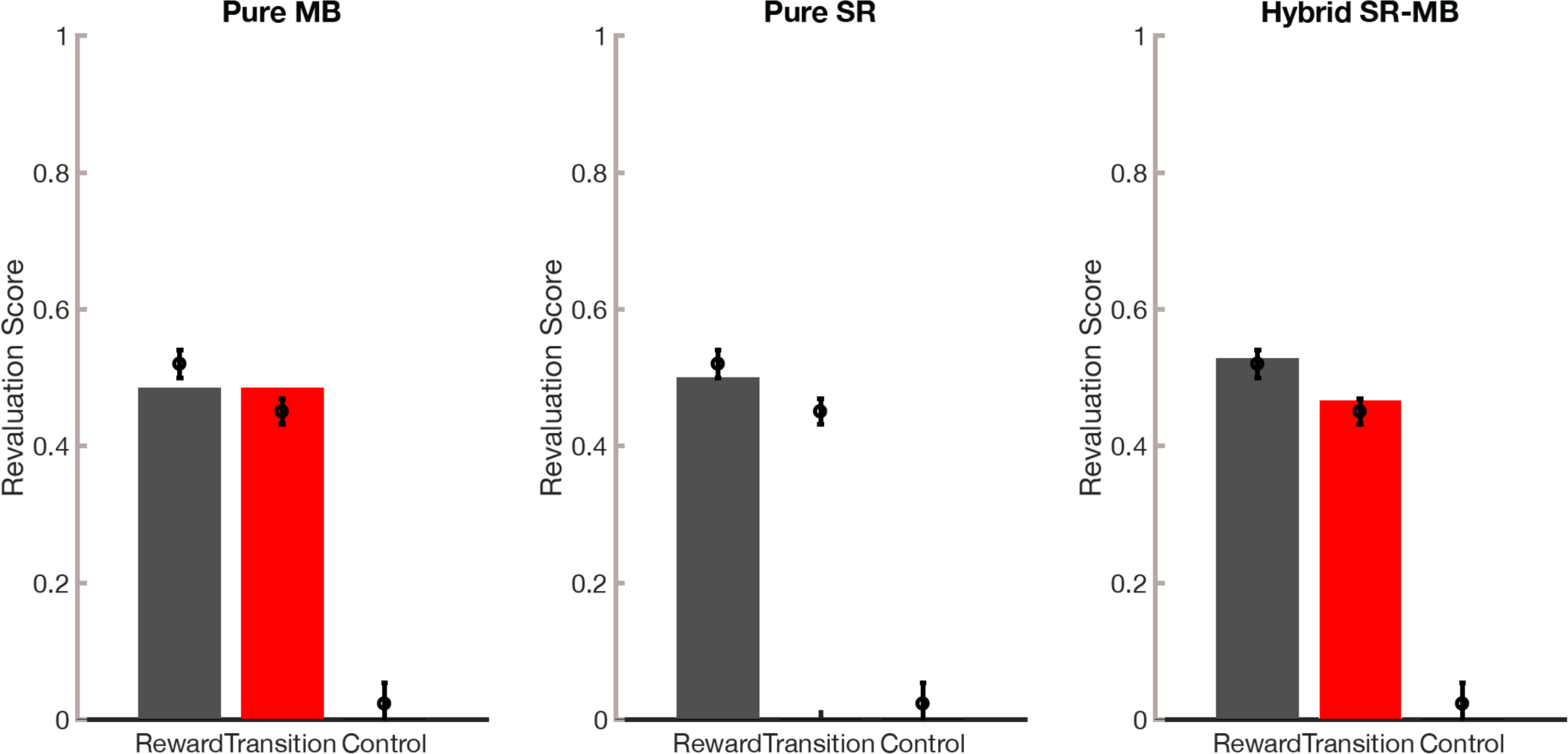
Model fits to the phase-3 test data from the passive learning task. We compared model performance against human data (black error bars) using a pure model-based learner (left), a pure SR learner (middle), and a hybrid SRMB learner (right). Human experimental results are represented in black and model performance is depicted in dark grey for reward revaluation, red for transition revaluation, and light grey for control. Human behavior is best explained by the hybrid account.

We did not hypothesize, nor do our results suggest, total reliance on an SR strategy; instead, we sought to investigate whether such a strategy is used at all by humans. A pure SR account does not by itself explain our data, because it predicts complete insensitivity to transition revaluation. In contrast, we see significantly greater revaluation in the transition revaluation condition compared to the control condition. This can be understood in terms of a hybrid SR-MB account analogous to the MB-MF hybrids^13,21^ considered previously (Figure 5, right panel). Although (as we discuss later) there are several ways to realize such a hybrid, we chose for simplicity to linearly combine the ratings from MB and SR algorithms. This linear combination allows the hybrid model to show partial sensitivity to transition revaluation. The hybrid model may also provide insight into the response time differences; under the assumption that effortful MB re-computation is invoked preferentially following transition revaluation (when it is, in fact, most needed), this condition would slow response time, consistent with our findings. Crucially, prior MB-MF hybrids^13,21^ cannot explain the asymmetry between transition and reward revaluation conditions (Figure 2C); among the theories we considered, this asymmetry is uniquely explained by a hybrid SR-MB account.

### Experiment 2: Differential sensitivity to revaluation types in a sequential decision task

In a second experiment, we sought to replicate and extend our results in two ways. First, Experiment 1 used a passive learning task, in which participants were exposed to a sequence of images, and the dependent measure was relative preference rating between different starting states. This is similar to previous Pavlovian experiments such as sensory preconditioning^,23^. Since a key purpose of state evaluation is guiding action choice, we sought to examine the same questions in the terms of decisions in a multistep instrumental task. This framing also allowed us to include an additional condition, which we call “policy revaluation.” The SR-based algorithms predict the same patterns of behavior in this condition as to the transition revaluation, but the actual sequence of participants’ experiences are much more closely matched to the reward revaluation condition. In particular, this condition turns on a change in reward amounts rather than state transition contingencies during the phase 2 relearning; no changes in the transition function occur in policy revaluation.

Participants completed four games, each of which corresponded to a different experimental condition (Figure 6). In each trial of each game, participants navigated through a three-stage decision tree (represented as rooms in a castle; see *Methods* for experiment details, Supplementary Figure 2). From the first stage (State 1), participants made a choice that took them deterministically to one of two second-stage states (States 2 and 3). Each second-stage state contained two available actions (and one unavailable action), and each action led deterministically to one of 3 reward-containing terminal states.

**Figure 6.**
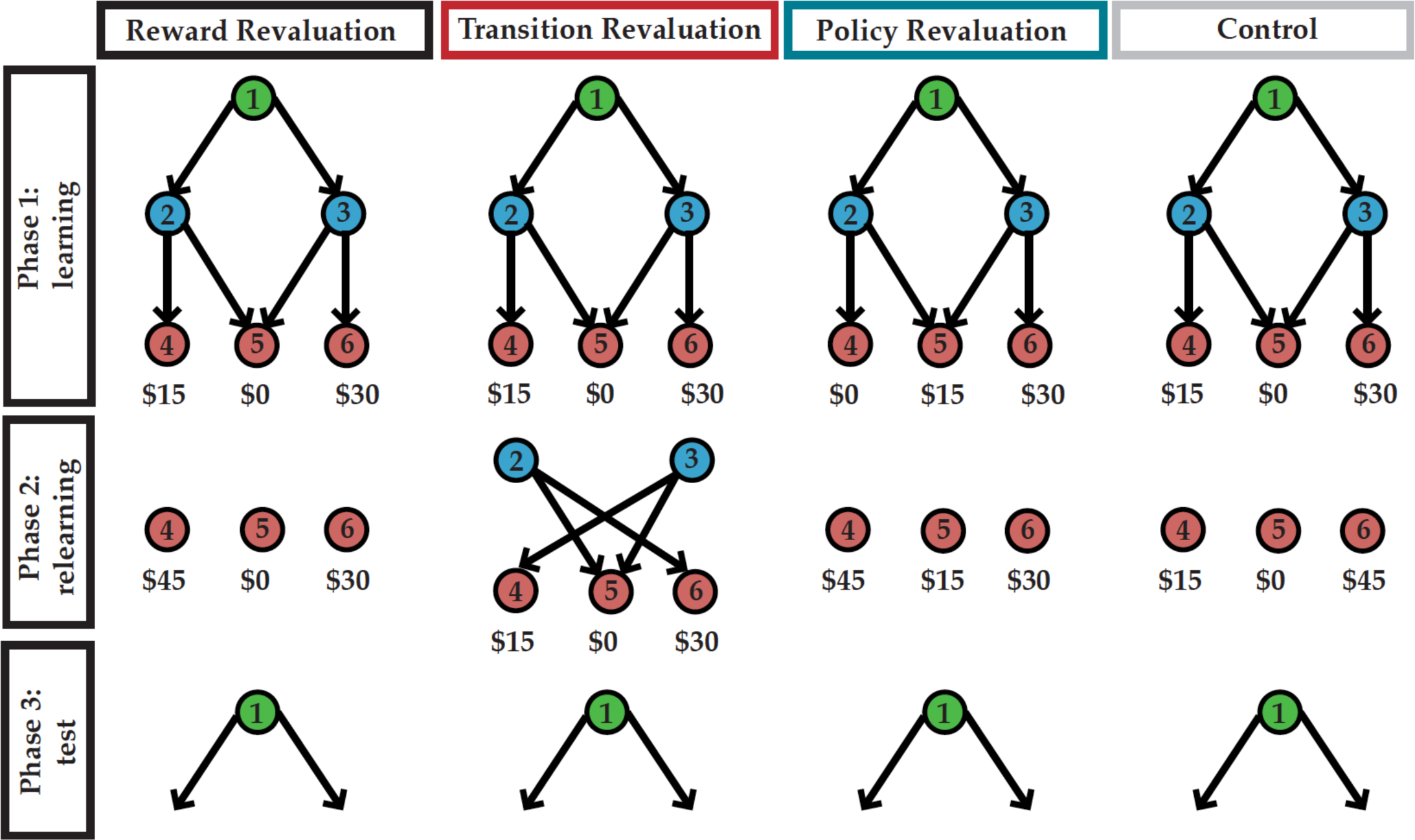
Schematic of the active sequential learning task. The underlying structure of each condition in Study II is represented in graphs. Numbered circles denote different states (rooms in a castle), and edges denote unidirectional actions available upon entering that state and the deterministic transition associated with those actions (that flow always from states with lower numbers to higher numbers, top to bottom in the schematic). Unavailable actions in States 2 and 3 are not shown. On each trial, participants were placed in 1 of the 6 states (castle rooms) and were required to make choices between upcoming states until they arrived in a terminal state and collected its reward. For a given phase of a given condition, trials began only in the states that are displayed in the figure for that condition and phase. For example, trials in Phase 2 of Reward revaluation condition began only from states 4, 5 and 6. In all conditions, State 1 contained 2 actions, both of which were always available. States 2 and 3 each contained 3 actions, however at any given time only 2 were available. Upon arriving in either state 2 or 3, the participant observed which actions were available and which were unavailable. For each condition, we measured whether participants changed their State 1 action choice between the end of Phase 1 and the single probe trial in Phase 3 from the action leading to State 3 to the action leading to State 2.

As in the previous experiment, these trials were grouped into 3 phases for each game (Figure 6). In Phase 1 (the learning phase), participants were trained on a specific reward and transition structure. If, for any condition, participants failed to perform the correct action from each non-terminal state on three of their last four visits to that state during Phase 1, they were removed from analysis. In Phase 2, (the relearning phase where revaluation could happen) a change in either the reward structure or the set of available actions occurred (the latter causing a change in the state-action-state transition function). Participants learned about the changed structure in 9 trials, such that they were exposed to the change at least 3 times. Importantly, as in Experiment 1, participants did not revisit the starting state in Phase 2, and hence never experienced any of the new contingencies following an action taken from the starting state. In Phase 3, participants performed a single test trial beginning from the starting state. For each condition, we defined the revaluation score as a binary variable indicating whether a participant switched their action in State 1 between the end of Phase 1 and the single probe trial in Phase 3.

**Figure 7.**
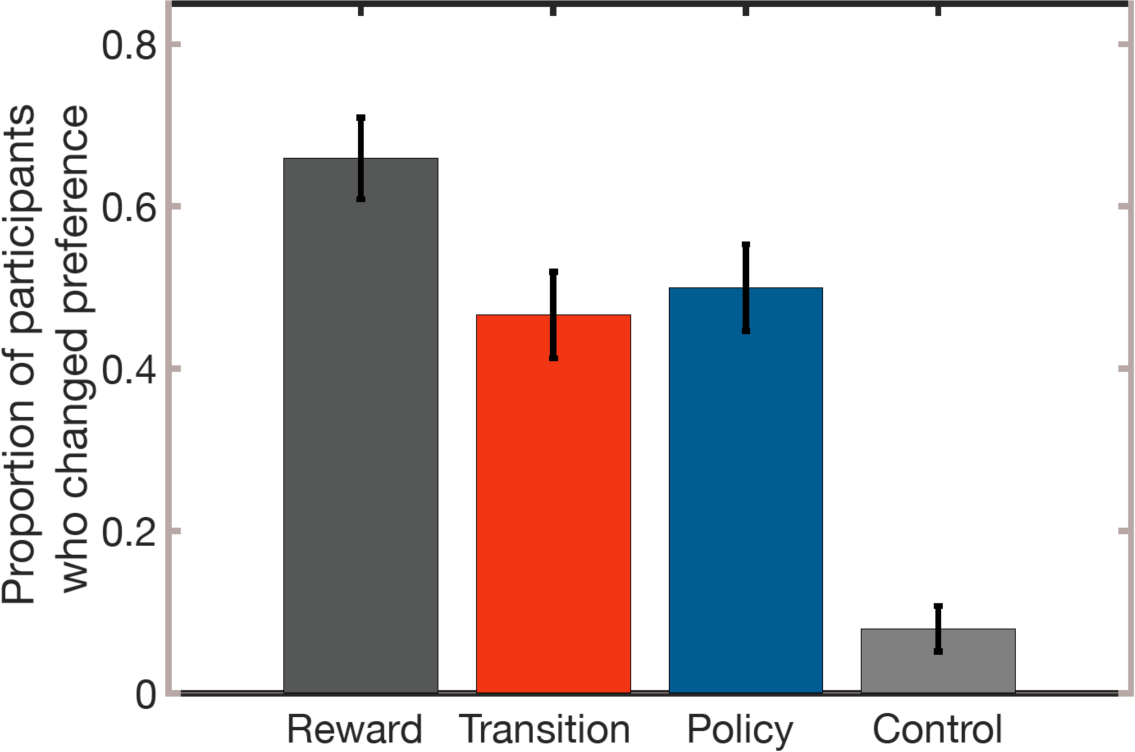
Behavioral performance in a sequential decision task. Proportion of participants (n=88) who changed preference following the relearning phase for reward, transition, and policy revaluation as well as the no revaluation control condition. Error bars represent 1 standard error of the proportion estimate.

Our results replicate those from Experiment 1, extending them to a new policy revaluation condition. The proportion of changed choices in the Phase 3 test, by condition, is shown in Figure 7 (Reward Revaluation: *mean = .6591, SEM = .0505*, Transition Revaluation: *mean = .4659, SEM = .0532*, Policy Revaluation: *mean = .5000, SEM = .0533*, Control: *mean = .0795, SEM = .0288*). Logistic regression verified that more participants successfully switched their stage-one action choice following the reward revaluation than the transition revaluation (*Contrast estimate = -.7958, Wald Z =-2.85, p = .0034*) and also the policy revaluation (*Contrast estimate = -.6592, Wald Z = -2.61, p = 0.0043*). In contrast, there was no significant difference between the proportions of participants that changed preference following policy revaluation compared to transition revaluation conditions (*Contrast estimate = .14, Wald Z = 0.56, p = 0.5771*). All three of the revaluation manipulations produced more switching than the control, no-revaluation condition (*reward > control contrast estimate = 3.1078, Wald Z = 6.95, p < 0.0001; transition > control contrast estimate = 2.31, Wald Z = 5.11, P < 0.0001; policy > control contrast estimate = 2.45, Wald Z = 5.20, p < 0.0001*), verifying that these results were due to a shift in preferences rather than nonspecific effects like forgetting. There was no significant effect of time on task (trial number) on revaluation score (F(1,190.6), P = .076). There was also no significant interaction of time on task with revaluation condition (F(2,69.49), P = 0.367).

### A hybrid model explains lower sensitivity to policy revaluation

The logic of the policy revaluation (Figure 6) is that the introduction of a large new reward at state 4 in the relearning phase should cause a change in the preferred action at State 2. The effect of this is to change which terminal state can be expected to follow the top-stage action that leads to State 2. Like transition revaluation, this manipulation should produce a change in top-stage preferences due to a change in the terminal state transition expectancies, but crucially it does so due to learning about reward amounts rather than the actual transition links in the graph. Because the SR caches predictions about which terminal state follows either State 1 action, it cannot update its decision policy without experiencing the newly preferred state along a trajectory initiated by the State 1 action leading to State 2. The MB and MF models (and the various hybrids) also treat this condition the same as the transition revaluation: in particular, the MB model should correctly re-compute the new Stage 1 action choice given learning about the new reward, whereas the MF model’s Stage 1 preferences should be blind to the change.

**Figure 8.**
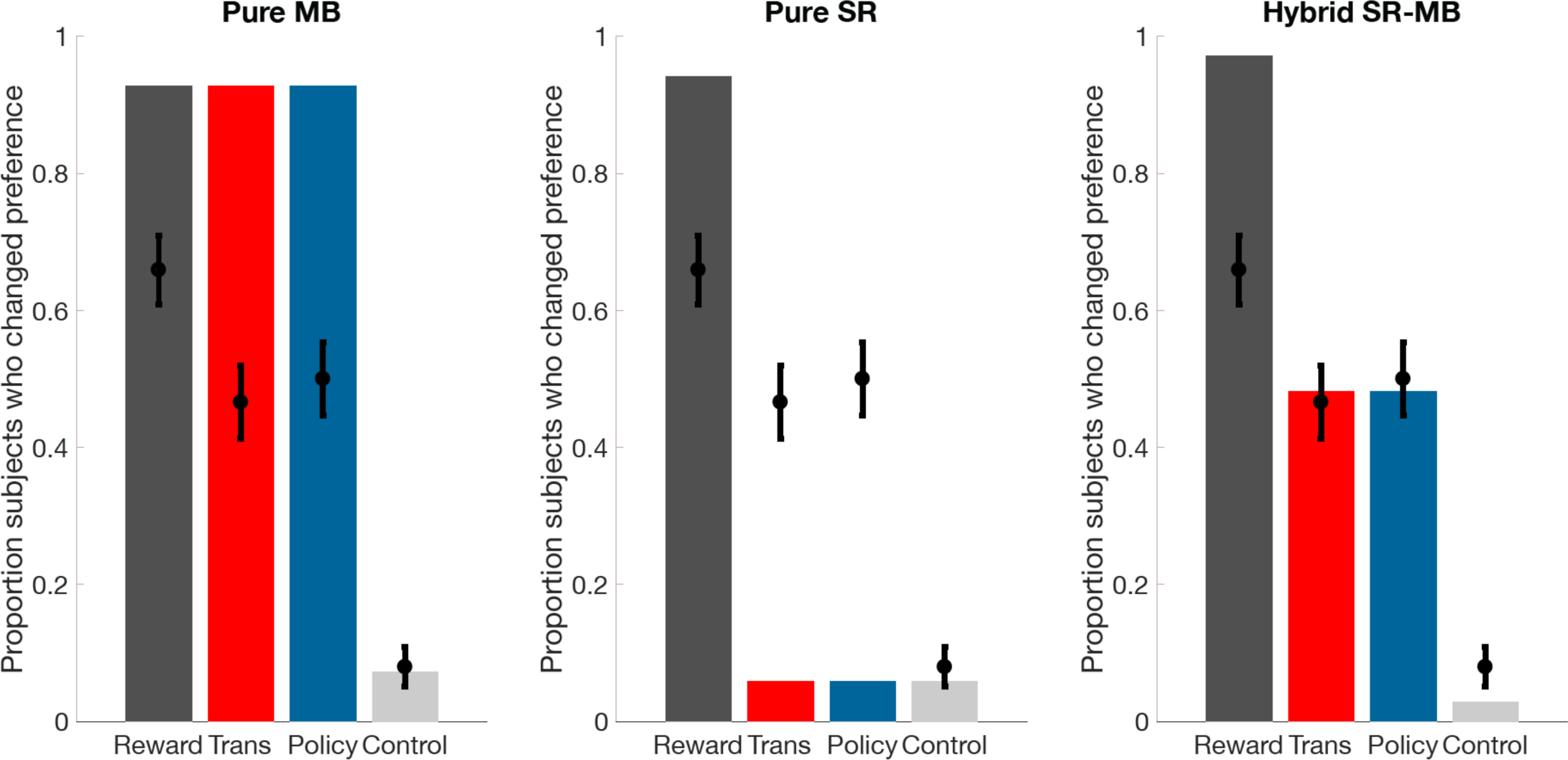
Model fits to the data from the sequential decision task. Behavioral data are represented with black error bars and model performance is depicted in grey for reward revaluation, red for transition revaluation, blue for policy revaluation, and light grey for control. Proportion of switches predicted by a pure model-based learner, a pure SR learner, and the hybrid SR-MB algorithm are shown for reward, transition, and policy revaluation conditions, compared to control (with matched learning rates for reward and transition learning). Consistent with results from the passive learning task, an algorithm using hybrid representations best captures human behavior. The X axis identifies the condition, while the y axis is based on proportions of participants who changed their preference.

The similarity in performance between the transition and policy revaluations suggests that the difference between transition revaluation and reward revaluation (here, and by extension, in the previous experiment) cannot be explained due to differences in acquiring the Phase 2 revaluation learning itself, about transitions than rewards. (For instance, consider an MB learner with a much slower learning rate for the transition matrix T than the learning rate R.) This is because the policy revaluation fools the SR in the same way as transition revaluation, but it does so by requiring participants to learn about a change in rewards rather than transitions. If participants were using a pure MB algorithm but were differentially skilled at transition and reward learning, we would expect policy revaluation to look more like reward revaluation than like transition revaluation, which was not the case.

We developed action-based variants of the models described in the previous section and fit them to the behavioral data (see *Methods* for details). Consistent with the results from the passive learning task, only the hybrid SR-MB model was able to adequately capture the pattern of differential sensitivity across conditions (Figure 8).

## Testing alternative possibilities

### Possibility 1: difference in MF-MB arbitration between conditions

It has been suggested that the relative balance between “state prediction errors” and “reward prediction errors” may be used for arbitration in a MB-MF hybrid learner^24^. In the policy revaluation condition, participants experience greater reward prediction error than in the reward revaluation condition, and thus the arbitration account would predict that participants should be more model-based and thus more successful at revaluation on policy revaluation trials compared with reward revaluation trials. This is the opposite of the SR prediction, and contradicted by our experimental observations.

To examine strategy changes further, we analyzed whether participants change strategy over the course of the experiment. Because only test trials can be used to ascertain strategy, and there is only one test trial per block, we cannot determine whether participants’ strategy changes within the course of a single block. If, however, participants’ strategy changed between blocks, over the course of the task, we would expect their revaluation performance to change as well. We thus used a repeated measures ANOVA to investigate whether there was an effect of accumulated time on the task (trial number) on revaluation score. Because a change in model choice would only affect revaluation scores of non-control trials, we eliminated control trials from this analysis. For experiment 1, this analysis revealed a significant effect of time on task on revaluation score [F(1, 57.259) = 9.9171, P < 0.01] indicating that participants improved at the task over time. If such a change in strategy over the task were responsible for the difference in revaluation score between reward and transition revaluation conditions, we would expect the effect of trial number on revaluation score to interact with the revaluation condition. However, there was no significant interaction of this effect with revaluation condition [F(1, 68.284) = .15436, P = 0.695]. For experiment 2 there was no significant effect of time on task (trial number) on revaluation score (F(1,190.6), P = .076). There was also no significant interaction of time on task with revaluation condition (F(2,69.49), P = 0.367). Thus, time on task cannot explain the difference between conditions.

### Possibility 2: Differences in learning and updating T, M, and R

For simplicity, we have assumed that during the relearning phase, the MB learner fully updates the experienced transitions in the transition matrix (T) and the SR learner fully updates the successor matrix M. However, performance on reward revaluation is not perfect. Could this be due to different learning rates for R and T? The relearning phase of experiment 1 included a probe trial once every 5 trials in which participants were required to choose between the second-level state in either sequence. Participants were unable to progress to the test phase until they chose the correct (highest value) state three times in a row. A repeated measures ANOVA was conducted to investigate the effect of revaluation condition (limited to reward revaluation trials and transition revaluation trials) on the number of trials required to meet the criterion for moving onto the test phase. There was no significant effect of revaluation condition: F(1,924) = 0.549, P = 0.359. Thus, we found no evidence for learning rate differences between conditions. Furthermore, results from policy revaluation conditions in experiment 2 shows that the findings are not merely limited to a difference between reward vs. transition learning, since in policy revaluation there are no changes in the transition probabilities (not updating of T or SR) and yet the behavior is more similar to transition revaluation rather than reward revaluation. Taken together, these findings are consistent with the idea that the differences between conditions are not merely due to differences in reward learning vs. transition learning.

## Discussion

The brain must trade off the computational costs of solving complex, dynamic decision tasks against the costs of making suboptimal decisions due to employing computational shortcuts. It has, accordingly, been argued that compared to model-based (MB) solutions, simple model-free (MF) learning saves time and computation at decision time at the cost of occasionally producing maladaptive choices in particular circumstances, such as rats working for devalued food. Here we consider a third strategy based on the successor representation (SR), which is noteworthy for two reasons. First, the SR caches temporal abstractions of future states. At decision time, while MB relies on forward search to evaluate actions, the SR simply retrieves cached representations of successor states and produces rapid, flexible behavior in many circumstances previously taken as signatures of the more costly MB deliberation. Second, the SR predicts (and our experiments confirmed) a novel asymmetric pattern of errors across different types of revaluation tasks. While MB performs equally well on all revaluation tests and MF solves none, the SR can use its cached representations to readily solve reward revaluation, but not transition or policy revaluation.

Previously, revaluation tasks – mostly reward revaluation – have been useful in distinguishing MB from MF predictions. However, MB and SR-based algorithms make similar predictions for standard reward revaluation tasks, which account for the bulk of evidence previously argued to support MB learning. By exploring other variants of revaluation (transition and policy revaluation), we were able to provide a direct empirical support for SR-based algorithms in human behavior. The crucial prediction made by the SR account, confirmed in two experiments, was that human participants would be more sensitive to changes in reward structure than to changes in transition and policy structures. Notably, even in absence of any changes in the transition structure in the policy revaluation condition, Experiment 2 showed that participants were also less sensitive to a shift in the optimal policy at intermediate states compared to the reward revaluation condition. This is consistent with SR-based algorithms but inconsistent with either MB algorithms or accounts of different MF-MB arbitration strategies^24^ following reward vs. state prediction errors.

It is important to stress that the SR is only one of a number of candidates for exact or approximate value computation mechanisms, and our study aimed to find affirmative evidence for its use rather than to argue that it can explain all choice behavior on its own. Studies using tasks with detour and shortcut manipulations^25^, particularly in the spatial domain, are conceptually similar to our transition revaluation. As in our study, some previous research suggests that organisms can in some circumstances also solve these tasks^26^. These results (together with more explicit evidence for step-by-step planning in tasks like chess or in evaluating truly novel compound concepts like tea jelly^27,2^) suggest some residual role for fully MB computation – or alternatively, that the brain employs additional mechanisms, such as replay-based learning, that would achieve the same effect^20^.

To reiterate, then, although our findings argue against a pure MB account (which would handle all our revaluation conditions with equal ease, or symmetrically), they also argue against a pure SR account, which predicts complete insensitivity to transition and policy revaluation (See Figures 2 and 8). Our data shows that people display significant revaluation behavior even in these conditions, though less than in the reward revaluation condition. Such results are expected under a hybrid SR-MB model in which decision policies reflect a combination of value estimates from MB and SR. We demonstrate that this hybrid theory provided a close fit to our data. It is best to think of the combination as a rough proxy for multi-system interactions, which are probably more complex^23^ than what we have sketched here. For instance, although we did not formally include or estimate purely MF learning in our modeling here, this is only because it predicts equally bad performance across all of our experimental revaluation conditions. We do not mean to deny the substantial evidence in favor of MF learning in certain circumstances, such as after overtraining. Indeed, MF learning may contribute to our finding that participants do not achieve 100% revaluation performance in any of our conditions, accounting for the slight difference between unnecessary switching in the control condition (which should measure nonspecific sloppiness like forgetting or choice randomness) and failure to fully adjust in the reward revaluation condition (see Figures 3 and 6).

Insofar as our results suggest that participants rely on a number of different evaluation strategies, they highlight the question of how the brain determines when to rely on each strategy (an arbitration problem). One general possibility is that humans use a form of meta-decision making, weighing the costs and benefits of extra deliberation to determine when to invoke MB computation^28,29^,30. This basic approach might fruitfully be extended to MB vs. SR as well as MB vs. MF arbitration. A meta-rational agent would be expected mostly to use the computationally cheap SR for flexible, goal-directed behavior (or even simpler MF for automaticity in stable environments), but would sometimes employ the more computationally intensive MB strategy to correct the SR-based estimate when needed (e.g., when transition structure changes). Given finite computational resources (and the problem that perfectly recognizing the circumstances when MB is required is potentially as hard as MB planning itself) this correction could be insufficient, leaving a residual trace of the biases induced by the SR. Our results on reaction times in the first experiment may provide a hint of such a hybrid strategy, since the MB system should take longer and might be more likely invoked in the transition revaluation condition (where it is, actually, needed).

Another form of SR hybrid could be realized by using the MB system (a cognitive map), or episodic memory replay, as a simulator to generate data for training the SR. This resembles the family of Dyna algorithms^20^. Evidence from rodents and human studies showing that offline replay of sequences during rest and sleep enhances memory consolidation^31^ and learning new trajectories^32,33^. Because the SR is updated via the simulations of the MB system or episodic memory offline, this Dyna-like hybrid model retains the SR’s advantage of fast action evaluation at decision time (Figure 8). Updating predictive representations via replay is in line with recent attention to the role of memory systems in planning and decision making^23,34^. These different realizations of an SR-MB hybrid are essentially speculative in the absence of direct evidence. Further work will be required to adjudicate between them.

All these models highlight the fact that the SR is itself a sort of world model, not entirely unlike the sorts of cognitive maps usually associated with hippocampus. The learnt representation is a predictive model, which allows mentally simulating distal future events rapidly, at least in the aggregate. It differs from the one-step model representations learned and used in standard MB learning, mainly because it aggregates these predictions over many future time-steps. This aggregation introduces a new free parameter: the timescale over which future events are aggregated. In the theory, the prediction timescale (known as the “planning horizon”) is controlled by the discount factor over future state occupancies in Equation 1, and need not, in general be the same as the agent’s time discount preference over delayed rewards^35^. Instead, we predict (and leave to future work to investigate) that the planning horizon should rationally be influenced by the statistical structure of experience, such as the stability or volatility of transitions and rewards in the environment. In other words, the structures of the environment should be reflected in the representations that are learned and stored in memory^40^. For instance, in more stable environments, it may be rational to cache representations with multistep contingencies over longer planning horizons, compared to volatile environments, where transition contingencies change frequently. In the unstable case, it would be counterproductive to cache contingencies beyond the hazard rate of the environment. This idea expands upon previous suggestion that environmental volatility should influence the use of MB revaluation vs. MF reward caching^36^. A further possibility, which remains to be tested, is that the brain might learn models at multiple timescales simultaneously (or build them up using offline replay) and later adaptively use representations for flexible planning at different scales^37^–39.

The SR hypothesis generates clear predictions about the neural representations underlying varieties of revaluation behavior, which could be tested in future functional neuroimaging studies. At least two major brain structures may underlie the SR: the medial temporal lobe (in particular, the hippocampus) and the prefrontal cortex. The hippocampus is implicated in the representation of both spatial^41^ and non-spatial^42,43^ cognitive maps^12^ (consistent with Tolman’s classic notion^8^), predictive representations of prospective goals^46^, as well as associative^47^, sequential^48^ and statistical learning^12,49^. Hippocampal replay processes help capture the topological structure of novel environments^33^ and sequentially simulate and construct path to future goals^50^ – beyond the animal’s direct experience – via forward and reverse replay^51^. This is consistent with recent fMRI and neural network modeling suggesting a potential role for the SR in complimentary learning systems, especially in the medial PFC and the hippocampus^12,52^. A recent modeling study^11^ suggested that the SR could explain the underlying design principles of place cells as studied in rodent electrophysiology. Taken together, these findings lend evidence to the hypothesis that the hippocampus may be involved in building and updating representation of SR’s predictive maps.

The second brain structure that may underlie predictive representations is the prefrontal cortex (PFC). A number of human studies have demonstrated the PFC’s role in the representation of prospective goals^53,54^. Lesions to the rat prefrontal cortex impair learning of transition structures (contingencies) but not incentive learning^26^. Ventromedial PFC is well connected to the hippocampus^47^ and is thought to mediate sampling information from episodic memory with the goal of decision-making^55^ and consolidation^56,57^, as well as the comparison and integration of value, abstract state-based inference^58^, and latent causes^59^. Furthermore, it has been suggested that the orbitofrontal cortex (OFC), may also be involved in ‘cognitive map’-like representations of task spaces^44^ and state spaces^45,60^. A recent finding suggests that the ventromedial PFC and the hippocampus encode proximity to a goal-state^61^. Together, the hippocampus and the OFC may be involved in forming and updating the SR, i.e. a rough predictive map of multi-step state transitions, according to simulated experience. Optimal decision-making may rely on the integration of OFC/ventromedial PFC and hippocampal cognitive maps consistent with our proposed hypothesis of hybrid predictive representations involved in decision-making. Testing the specific role of the prefrontal and hippocampal contributions to the successor representations offers an exciting avenue for future functional neuroimaging studies.

In short, we have shown that human behavior reveals the contribution of a particular sort of internal model of outcome predictions, the successor representation (SR). We designed varieties of revaluation tasks in which different algorithms for representation learning predict different planning and decision-making behavior. In contrast to learning only one-step representations, as in classic model-based learning, the successor representation stores multi-step predictive representations of future states. These predictive representations can be learned via mechanisms such as temporal difference learning, and can be updated via multiple routes: via direct experience, via interaction with rolled out model-based predictions, or via simulated experience or offline replay. We have shown that human behavior under different varieties of revaluation reveals the contribution of such predictive representations. Future studies can explore the individual differences observed here to study the flexibility and pathologies in arbitration of different representation learning approaches in planning and decision-making. We anticipate these findings to open up avenues for computational, electrophysiology, and neuroimaging studies investigating the neural underpinnings for this evaluation mechanism.

## Methods

### Task 1: Sequential learning task

69 (*mean age = 22.2, STD = 4.6, 42 female*) participants were recruited for the passive learning task, of which 4 participants were excluded as they did not learn the task and could not finish the study within the allotted 1.5 hours. 7 were removed from final analysis due to accuracies below 80% in the categorization task (described below), a threshold used as a measure of attention to (engagement with) the experiment, leaving 58 participants. The study design and the collection of data complied with all relevant ethical regulations. Princeton University’s ethics committee approved the study. All participants signed an informed consent form, reported no history of mental illness, and had normal or corrected to normal vision.

Participants played 20 games, each corresponding to one of three conditions: reward devaluation (8 games), transition devaluation (8 games), and a control condition (4 games). Each game had three phases: 1) a learning phase, 2) a revaluation phase, and 3) a test phase. Games of various conditions were randomly interleaved for each participant. In Figure 2, the schematic of all phases and two experimental conditions (reward and transition revaluation) are shown as state transition diagrams. Here ‘states’ are represented as numbered circles and arrows specify one-step deterministic transitions. Each state was uniquely tagged in each game with a distinct image (of either a face, scene, or an object, Figure 2). The stage of the current state within a multi-stage trajectory was indicated by the distinct background color of that state (e.g., state 1 had a green background, state 2 blue, and state 3 red; Figure 2, Supplementary Figure 1).

During phase 1 (learning phase) of each game, participants first experienced all states and their associated reward. They passively traversed 6 states and learned the transition structure that divided them up into 2 trajectories. To ensure that participants attended to each state, participants were asked to perform a category judgment on the images associated with each state (face, scene, object). Phase 1 was concluded once the participant reached a learning criterion, which was reached if the participant preferred the middle state of the most rewarding trajectory (preference between state 3 vs. 4 in Figure 2, Supplementary Figure 1). The criterion was tested every 5 trials: participants were shown the middle states of each trajectory (blue background in Figure 2) of the two trajectories and asked which one they preferred. For each trajectory, learning phase criterion was reached once their preference indicated the middle state of the optimally rewarding trajectory, or after 20 stimulus presentations. Trials in which participants did not reach learning criterion within the allotted 20 stimulus presentations were excluded from further analysis. During the final test phase, participants were once again shown the starting state of the two trajectories and asked to indicate which one they preferred (i.e., which one led to greater reward) on a continuous scale.

During phase 2 (revaluation phase) participants passively viewed all states except the starting states of each trajectory (states 1 and 2 in Figure 1); trajectories were always initiated in one of the second-stage states. As in phase 1, participants performed a category judgment on the images of the states they visited. This category task served as a measure of attention to the states during both phases. In the control condition, there were no changes to the task structure. In the reward revaluation condition, the rewards associated with the terminal states of the trajectories were swapped. In the transition revaluation condition, the connectivity between the second- and third-stage states was altered, such that the middle state of a given trajectory now led to the final state of the other trajectory (Figure 2). As in phase 1, participants were probed for their preference of the middle states every 5 stimuli, and phase 2 concluded once they met the learning criterion (3 correct decisions about the middle states) or after 20 stimuli. During phase 3, participants were instructed to once again rate their preference for the start states.

### Experiment 1: Computational models

We compared the performance of three models to human behavior: a model-based learner that computes values using its knowledge of the transition and reward functions (Figure 4, left), a pure SR learner that computes values using estimates of the reward function and SR (Figure 4, middle), and a hybrid SR-MB model that linearly combines the ratings of the two learners (Figure 4, right).

The SR learner uses two structures to compute state value: a vector R (the reward function encoding the expected immediate reward in each state) and a matrix M (the expected discounted future occupancy in each state):

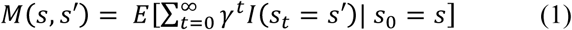

where γ is a discount parameter. Since in our task the terminal states are absorbing, we set γ=1 (i.e., no discounting). The SR learner combines these two structures to compute the value of a state by taking the inner product of R and the row of M corresponding to that state:

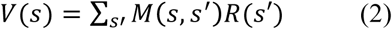

The model-based learner computes state values by iterating the Bellman equation over all states until convergence [cite Sutton & Barto]:

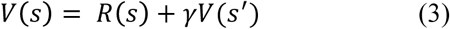

where *s*’ is the immediate successor of state *s*.

We assumed that preference ratings were generated by a scaled function of the state values:

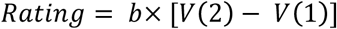

where b is a free parameter. For the SR-MB hybrid model, we assumed that the preference rating was a linear combination of the ratings generated by the two component models:

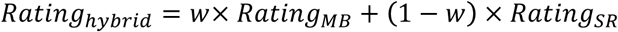

where *w* is a free parameter.

Note that our strategy here is to model such ratings predicted by the different algorithms’ representations, rather than the trial-by-trial learning process that produced these representations. This is because the structure of the task does not provide enough variability in participants’ experience, or monitoring as to participants’ ongoing beliefs, to constrain trial-by-trial learning within the acquisition phases. In particular, because the experienced rewards and transitions are deterministic within a given phase of each game, variables like learning rates, which would govern the rate at which model representations reach their asymptotic values, are under-constrained. Furthermore, the task is passive; participants’ beliefs are tested only sporadically and indirectly with relative preference judgments.

We thus assume that by the end of each phase, each model representation has reached its asymptotic value, consistent with the information presented and the experiences permitted during that phase, and with the usual learning rules for these algorithms. Specifically, we assume that at the end of phase 1 and again following phase 2, the model-based learner has appropriately updated the transition function (providing which *s*’ follows which *s*) to the most recently experienced contingencies, the SR learner has appropriately updated *M*(*s*, *s*’), and both learners have appropriately updated *R*(*S*). Importantly – to capture what would be the endpoint of trial-by-trial learning of the different representations – in each case we assume the various representations are only updated for all states *s* visited during a phase; representations for states not visited in phase 2 remain unchanged. Using these updated representations, we compute *V*(*s*) at the end of phase 1 and again end of phase 2 using equation 2 for the SR learner and 3 for MB learner. *V*(*s*) is used to derive ratings at the end of phase 1 and the beginning of phase 3. Revaluation scores are then computed by subtracting the phase 1 rating from phase 3 rating:

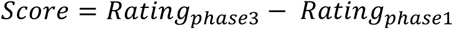

A subject’s revaluation score for a given trial was modeled as being drawn from a Gaussian distribution, centered on the model’s predicted score, for that revaluation condition:

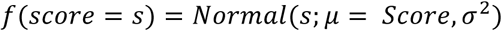

where f is the density of a score s. Here σ^2^ is free parameter.

Altogether, SR and MB each had two free parameters (*b*, σ^2^) and the hybrid model had three (*b*, *w*, σ^2^). For each participant, we estimated parameters *b* and *w*, by maximizing the likelihood of her revaluation scores, jointly with the group-level distributions over the entire population using an Expectation Maximization procedure^62^. For each subject, the maximum likelihood estimate of σ^2^ was directly computed based on the residuals between the model estimates (determined by the other parameters) and the data:

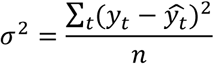

where y_t_ is the subject’s revaluation score on trial *t*, 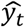 is the model’s predicted revaluation score for trial *t*, and *n* is the number of trials. In order to constrain *w* to be between 0 and 1, the likelihood function passed the input to parameter *w*, *w_in_* through the transformation 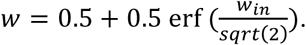

Aggregate log-likelihoods for each model were computed using a leave-one-out, subject-level, cross validation procedure. This involved, for a given subject *j*, using data from every other subject *y* ≠ *j* to fit a group level parameter distributions. The log likelihood of subject j’s ratings was then computed averaging the log-likelihood of 10,000 samples taken from the group distribution. We repeated this procedure for each subject and computed aggregate log-likelihood by summing over subjects.

### Experiment 2: Sequential decision task

This task was run on Amazon’s Mechanical Turk (AMT) using Psiturk software^63^. We set out to collect data from 100 participants, and 112 participants (*mean age = 33.6, STD = 10.5, 53 female*) were recruited to complete the experiment. All participants were required to achieve 100% accuracy on a 9-item instruction comprehension task before beginning the task. Participants were excluded for failing to learn the appropriate decision policy at the end of phase 1 training (Preference 1) in at least one of the conditions (see below). After collecting 100, we removed those who failed to pass the exclusion criteria and then continued to collect until we had an equal number of non-removed subjects in each of the four experimental conditions (*88 participants, mean age =33.8, STD = 10.8, 45 female*) each of which corresponded to an order of revaluation trial types. Participants received a bonus proportional to the total amount of reward collected.

Participants made choices in order to collect rewards by navigating an avatar through the rooms of a castle. The underlying structure of each condition of the task is displayed schematically in Figure 5. In each trial, participants were placed in 1 of the 6 states (castle rooms) and were required to make choices until they arrived to a terminal state and collected the associated reward. States were displayed as colored shapes on a screen. The spatial position and color of each state was randomized across blocks, yet remained fixed within a block.

Each participant performed four blocks of trials. Each block corresponded to a different condition. Block order was counterbalanced across participants according to a Latin square design. Each block consisted of 3 phases (Figure 5). In the learning phase (phase 1) participants were trained on a specific reward and transition structure. Training involved completing 39 trials. The starting state for each trial was randomized so that at least 14 trials began from State 1, at least 7 trials began from State 2 as well as State 3, and at least 2 trials began in each terminal state. In each condition, the reward and available actions for phase 1 were arranged so that State 6 contained the highest reward and was exclusively accessible from State 3. Thus, by the end of phase 1, participants should have learned to select the State 1 action leading to State 3. The other terminal states respectively contained, low and medium sized reward. One of the other terminal states was accessible from both States 2 and 3 and one was accessible exclusively from State 3. This arrangement ensured that there was a ‘correct’ action from each non-terminal state that would lead to a higher reward. If, for any condition, participants failed to perform the correct action from each non-terminal state on three of their last four visits to that state, they were removed from analysis.

In phase 2, a change in either the reward associated with one of the terminal states or in the set of available actions in States 2 and 3 occurred. In the reward revaluation condition, the amount of reward in State 4, which previously contained the highest reward accessible from State 2, was increased so that this state was now the most rewarding terminal state. This change thus altered the reward of the state that the participant had previously experienced as following the State 1 action leading to State 2. In the transition revaluation condition, the set of available actions in States 2 and 3 was changed so that State 6, the terminal state containing the highest reward could be reached exclusively from State 2. This change thus altered which terminal state could follow either State 1 action and was thus comparable to the transition revaluation in passive learning task. In the policy revaluation condition, the amount of reward in the terminal state containing the smallest reward was increased so that this state now contained the highest reward. Because this would alter which action was preferred from State 2, it changed which terminal state would be expected to follow the action leading to State 2. Thus despite involving a reward change, this change would have similar effects on successor representation models as the transition revaluation. Finally, in the control condition, the amount of reward in the state containing the highest amount of reward increased. In each condition, phase 2 consisted of 9 trials. In the reward revaluation, policy revaluation and control conditions, these trials started 3 times from each terminal state. Phase 2 trials in the transition revaluation started 3 times from both State 2 and State 3, so as to allow participants to observe the change in available actions, and 1 time from each terminal state. Crucially, participants did not visit the start state (state 1) during phase 2, and hence never experienced any changes in reward following an action taken from the start state. In the test phase (phase 3), participants performed a single trial beginning from the start state. In phase 3, participants completed a single trial starting in State 1. We defined the revaluation score as 1 if they switched to the now better action leading to State 2 and 0 if they stayed with the action leading to State 3.

### Logistic regression analysis

All of our descriptive analyses involved performing pair-wise comparisons between proportions of participants that switch action preference following different revaluation conditions. In order to perform such pairwise comparisons while correctly accounting for the repeated-measure structure of the experiment, we fit a logistic regression model where the dependent variable was a binary indicator of whether a given participant changed action preference in state 1 between phases 1 and 3. The model had four independent variables: a binary indicator variable for each condition that was set to 1 when the given response was from that condition. This model provided a coefficient estimate for each condition indicating the logit-transformed probability that participants switched state 1 action preference in phase 3 of that condition. To obtain standard errors on coefficient estimates that accounted for participant-level clustering due to the repeated measures, we employed a cluster-robust Huber-White estimator (using the robcov function from the R package rms^64^). Contrasts between coefficients were computed by fitting the model once for each condition, substituting that condition in as the intercept so that coefficient estimates for the other three conditions represented contrasts from it.

### Experiment 2: Computational models

**∊-greedy model:** All learners convert action values, *Q*, to choice probabilities using an ∊-greedy rule. This rule chooses the available action with the max *Q* value with probability 1 – *∊* and chooses a random available action with probability *∊*. Thus, for available actions *sa* in state *s*:

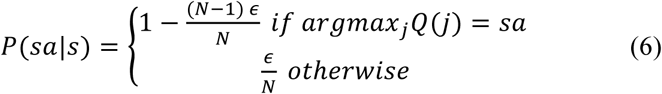

where N is the number of actions available in state s’. We consider terminal states to have a single available action. We set *P*(*sa*|*s*) = 0 for all actions not available in state *s*.

As in our modeling of the passive learning task, the SR learner uses two structures to compute value: a reward vector *R* and a matrix of expected future occupancies, *M*. The only change here is that the elements of *M* are indexed by actions, *sa*. Likewise, *R*(*sa*) stores the reward associated with taking action a from state *s*. *M*(*sa*, *s’a’*) stores the expected future (cumulative, discounted) number of times action *s’a’* will be performed on a trial following action *a* from state *s*, *sa*. The SR learner combines these two structures to compute the value of an action by taking the inner product of the reward vector R and the row of M that corresponds to that action in the current state:

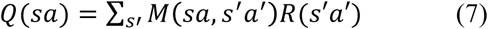

The model-based learner computes value estimates by combining its knowledge of the transition and reward structure, iterating the following Bellman equation until convergence:

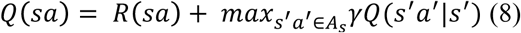

where *s*’ is the state to which *s’a’* transitions and *A_a_* is the set of actions, *s*, *a’*, avaialbe in state *s’*.

The SR-MB hybrid learner forms action probabilities by combining action probabilities from both SR and MB learners *P_sr_*(*sa*|*s*) and *P_mb_*(*sa*|*s*). The model assumes that the two action probabilities are combined according to a weighted average:

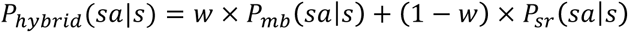

where *w* is a free parameter.

As with the passive learning task, because parameters like learning rates are under constrained, we assume that by the end of each phase, each model representation has reached its asymptotic value, appropriately updated according to the information presented and the experiences permitted during that phase. For the active learning task, this means that at the end of phase 1 and also phase 2, the MB learner has appropriately updated *A_s_*, the SR learner has updated *M*(*sa, s’a’*), and both learners have adjusted *R*(*sa*), but again in each case only for all states *s* visited and actions *sa* performed during that phase.

Using these updated representations, we compute *Q*(*s*, *a*) at the end of phase 1 and again end of phase 2 using equation 7 for the SR learner and 8 for MB learner. *Q*(*s*, *a*) is used to derive action for each action at the end of phase 1 and also at the beginning of phase 3.

As in experiment 1, we computed an aggregate log-likelihood of each subject’s phase 3 choice under each model, using a leave-one-out, subject-level cross validation procedure. Parameters *w* and *∊* were constrained to be between 0 and 1 by being passed through the same transformation used to constrain *w* in experiment 1. For each subject, we fit a distribution of group level parameters to subject’s choices, using data from every other subject. In order constrain noise parameters (*∊*), we included the subjects last four choices in phase 1 from each decision state in addition to the subject’s phase 3 choice in the likelihood function used to estimate parameters. We chose these choices because our exclusion criterion assumed that subjects learning would have reached asymptote by this point (and subjects were excluded if they did not perform the correct action in 3 of their last 4 visits to each of these states). We then compute the log likelihood of subject *j’s* phase 3 choices by averaging the log-likelihood of this choice using 10,000 samples taken from the group distribution. The aggregate log-likelihood of the phase 3 choices under the model was computed by summing over the individual likelihoods computed for each subject.

### Replication and randomization

While the two experiments are conceptual replications of one another, using different designs and choices approaches to compare similar conditions, we did not conduct individual replications of each experimental design. We employed within-subject designs: each participant underwent equal number of all conditions in a randomized fashion. The order of experimental conditions was randomized by the experiment’s code; therefore, investigators were blind to the specific order for each participant.

### Data and code availability

Source data (Figures 4 and 7) and costume code for generating the models (Figures 5 and 8) will be made available upon request.

## Acknowledgements

This project was made possible through grant support from the National Institutes of Health CRCNS award 1R01MH109177, the National Institutes of Health under Ruth L. Kirschstein National Research Service Award 1F31MH110111-01, and the John Templeton Foundation. The authors declare no competing interests. The authors would like to acknowledge Ken Norman and Ross Otto for helpful conversations, and Alex Rich and Shannon Tubridy for assistance with psiturk. The opinions expressed in this publication are those of the authors and do not necessarily reflect the views of the funding agencies. The funders had no role in study design, data collection and analysis, decision to publish or preparation of the manuscript.

## Supplementary Material

### The successor representation in human reinforcement learning

Momennejad I^1*^, Russek EM^2*^, Cheong JH^3^, Botvinick MM^4^, Daw N^1^, Gershman SJ^5^

**Supplementary Figure 1.**
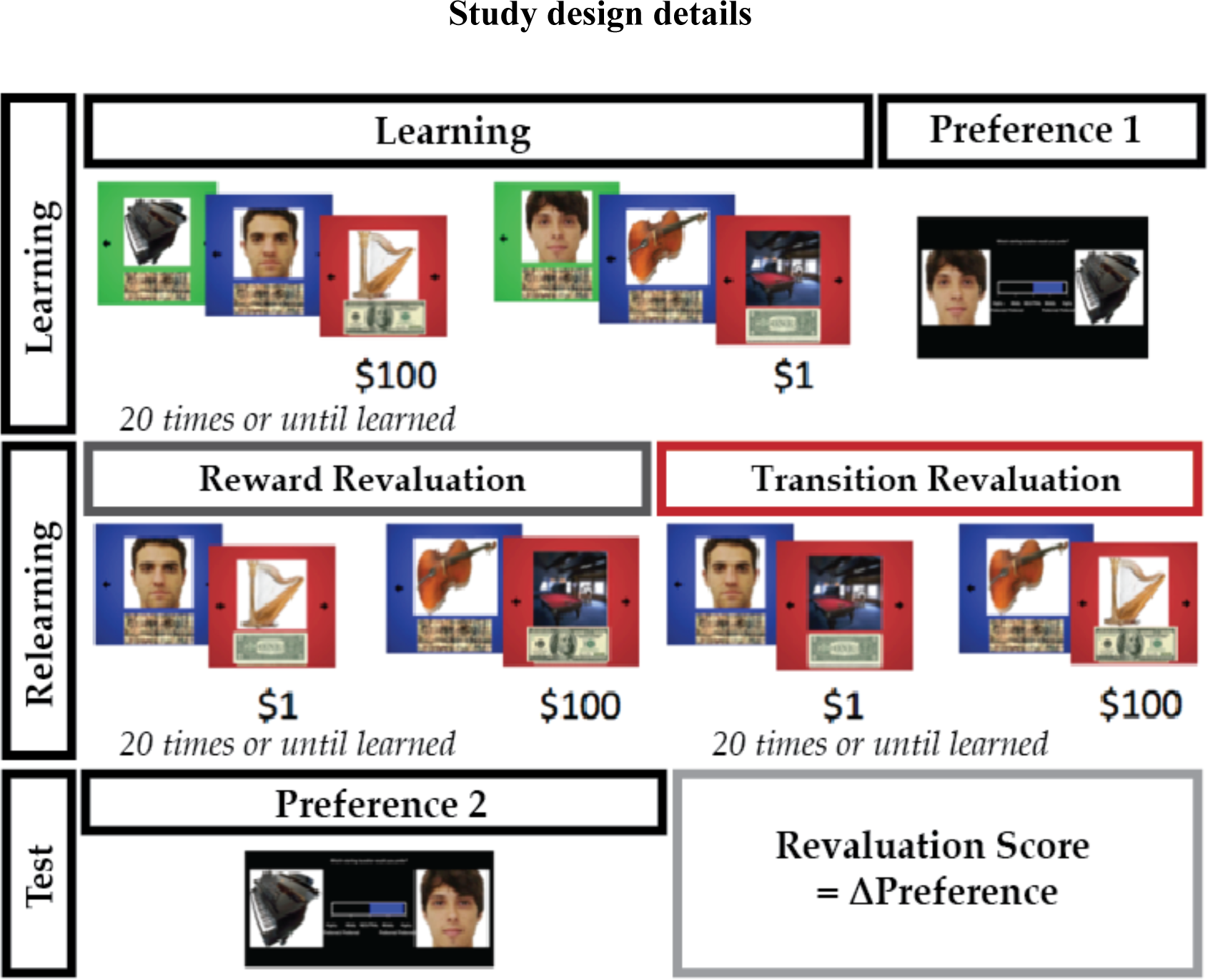
Detailed time course of stimuli for the first experiment are depicted. The cover story instructed participants that they were photographers that will take photos on a ride starting from 2 starting stations (green). Each trajectory started in a given station (green) then passed through a middle station (blue) and ended at a final station (red). As such, the color context always indicated where along a trajectory the participants were. Every station had a unique picture of a face, scene, or object: participants were told this is the photograph they took. Below each photograph appeared a scrambled dollar bill or a complete one, indicating how much the participant would earn by selling that photograph. The goal of the game was to indicate at the end preference for one of the starting states that participants had learned about, and the basis of this preference was to maximize the reward of the photos they took. In order to ensure that participants were paying attention to the unique images on screen, they were asked to perform a categorization task were they would indicate whether the object on screen was a face, a scene, or an object. Their performance on the category task served as an indication of how much attention they were paying to the categories at hand. During the learning phase participants visited the trajectories 20 times, or until they satisfied the learning criterion. The learning criterion was based on their performance on intermittent preference tests, where they were probed for their preference of the middle station (blue background) every 5 trials. This allowed us to know that they learned the associative value of the trajectories. After the learning phase, participants indicated their preference for either one of the starting states (green background) on a sliding scale (Preference 1 in the figure). Following Preference 1 ratings, participants entered the relearning phase, where they did not visit the starting state (green background) any longer, but only visited the middle and final states. During control trials, these transitions remained unchanged compared to the learning phase. In trials in the reward revaluation condition, participants experienced a change in rewards associated with the final states, and in trials in the transition revaluation condition they experienced a change in the transition from the middle states (stations with the blue background) to the final states (stations with the red background), as indicated in the Figure. As in the relearning phase, participants were probed for their preference of the middle stations after every exposure to 5 stimuli. This was used as a learning criterion, and participants were exposed to stimuli 20 times or until they satisfied learning criterion. After the relearning phase, participants entered the test phase (Preference 2) where they were shown the starting states (stations with the green background) and asked again to indicate their preference for one or the other using a sliding scale as in Preference 1. By subtracting the preference scores in Preference 2 from Preference 1, we get the revaluation score, which tells us whether a participant, on a given trial, has changed their preference. The direction or sign of the change in preference denotes whether participants revalued their initial preference.

**Supplementary Figure 2.**
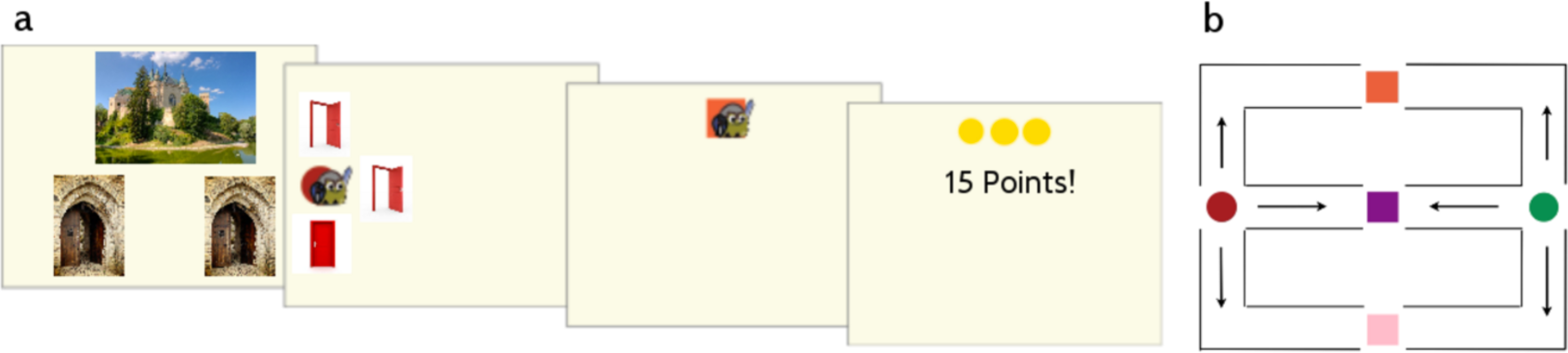
(a) Example trial from experiment 2, beginning in state 1. In state 1, the participant saw an image of the outside of the castle they were in and were required to select one of two entrances. Each condition had a unique castle image as well as unique doors that were constant for the trials in that condition. In this trial, the participant selected the left castle entrance. Selecting the left entrance transitioned the participant to the circle room on the left side of the castle and selecting the right entrance transitioned the participant to the castle room on the right. Upon arriving in the circle room, the participant observed three doors, two of which were open. Upon selecting one of the open doors, the participant transitioned to one of three-square rooms. Upon pressing ‘space bar’ in one of the square rooms, the participant received the reward contained in that room. (b) Transition structure within the castle. State 1 was outside the castle. States 2 and 3 were represented as circle rooms on the left and right side of the castle. Which side of the screen States 2 and 3 were on was randomized across blocks yet constant within a block. States 4, 5 and 6 were represented as squares that lied along a column middle of the screen. The position (top, middle or bottom) of States 4, 5 and 6 was randomized across blocks yet constant within a block. Similarly, the color of states 2-6 was randomized across blocks, yet constant within a block. Arrows denote transitions from States 2 and 3. For example, selecting the door above the circle in either state 2 or 3 always took the participant to the state represented by the square room on the top of the screen. Each door was open when the action it corresponded to was available.

## Supplementary methods and results

### Modeling with a soft-max choice rule

We additionally modeled experiment 2 using a softmax choice rule (Supplementary figure 3 and Supplementary Table 1). For this model, Q values for the SR and MB learners were formed the same as in the *∊*-greedy model. The SR and MB learners each converted action values, Q, to choice probabilities using a logistic soft-max, P(sa|s) ∝ exp (βQ(s, a)), normalized over all available a in state s. Here β is a free parameter. The SR-MB hybrid learner formed action probabilities by combining the Q values from both SR and MB learners: P_hybrid_(sa|s) ∝ exp(β_MB_Q_MB_(s,a) + β_SR_Q_SR_(s,a)), again normalized over all available a in state s. The SR and MB learners each contained one free parameter (β_MB_, β_SR_). Group-level models were fit and log-likelihood was computed in the same manner as for the ∊-greedy model.

In accordance with the ∊-greedy model, the softmax model also shows that a hybrid model, including the successor representation augmented with a model-based strategy (or offline training), is best able to explain the data. Qualitatively (as demonstrated by the simulations using the best fit parameters) only a model that includes the successor representation is capable of explaining the asymmetry in test performance following reward compared to transition and policy revaluation conditions.

**Supplementary Figure 3.**
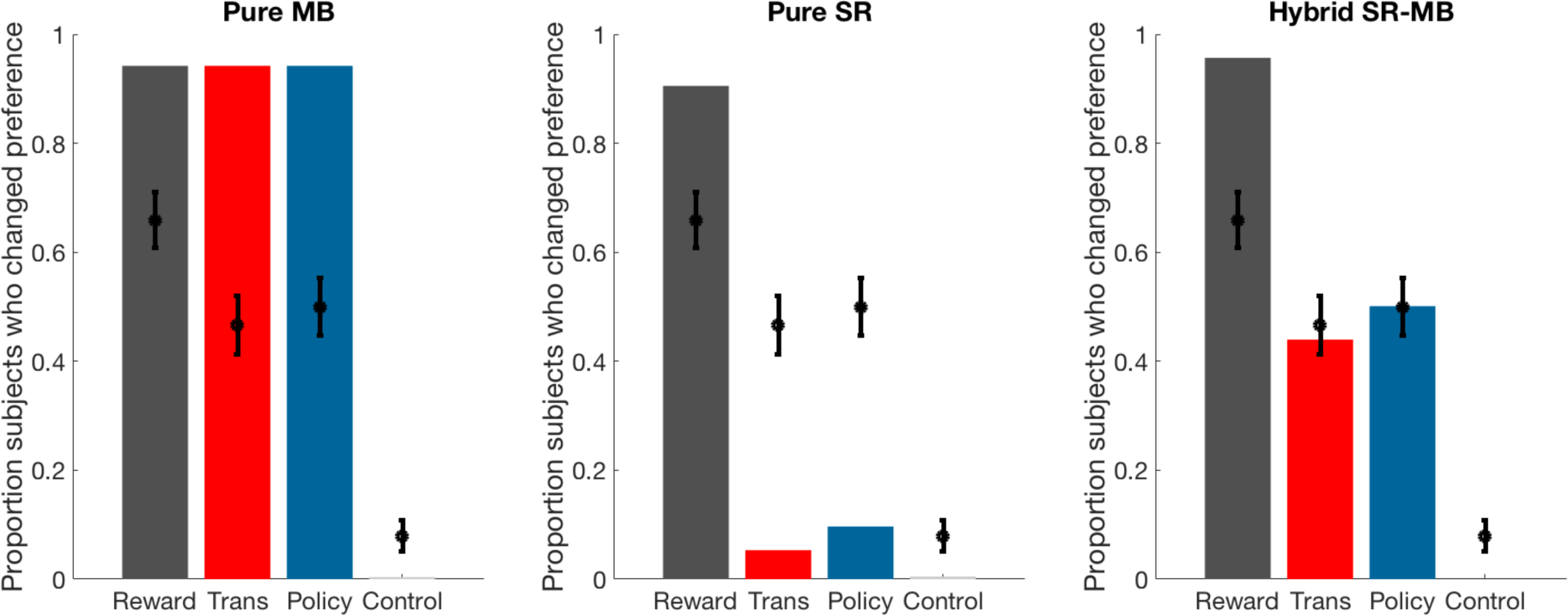
Soft-max model predictions (at best parameters for group fit).

We also report cross-validated model log-likelihoods, along with parameter fits in supplementary tables 1–3. Due to the relatively small number of choice trials, we do not view the experiments as being ideal for model comparison. Rather, we view the purpose of model fitting and simulation to illustrate the clear qualitative predictions made by the different models.

**Supplementary table 1.** Model fits for experiment 2, soft-max model

| | Bmb ($\mu$) | Bsr ($\mu$) | Log-likelihood |
| --- | --- | --- | --- |
| MB | 0.185796 | x | -378.4640787 |
| SR | x | 0.182304 | -343.6524022 |
| MB+SR | 0.103172 | 0.115476 | -248.828638 |

**Supplementary table 2.**
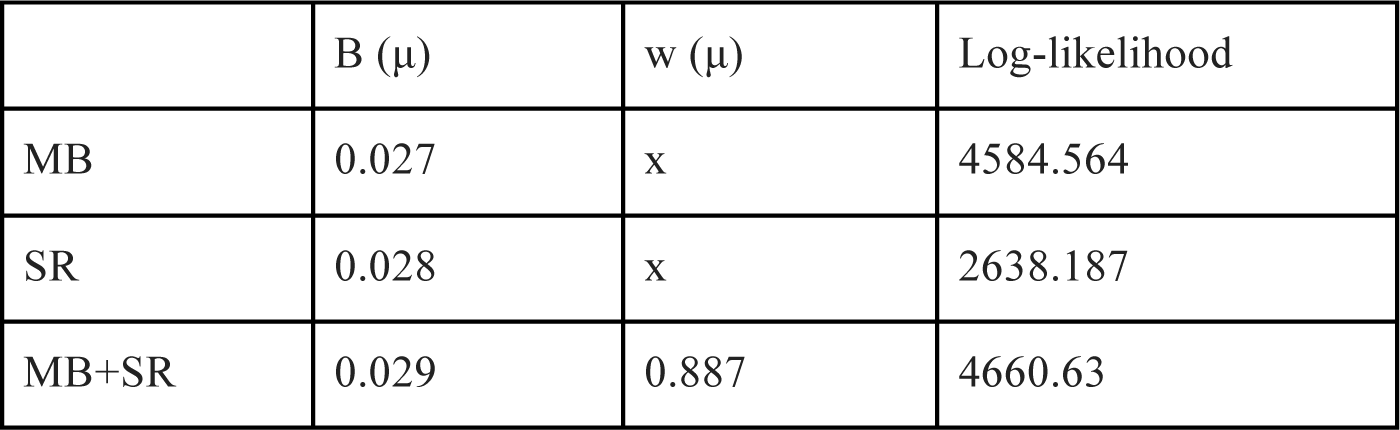
Model fits for experiment 1

| | B ($\mu$) | w ($\mu$) | Log-likelihood |
| --- | --- | --- | --- |
| MB | 0.027 | x | 4584.564 |
| SR | 0.028 | x | 2638.187 |
| MB+SR | 0.029 | 0.887 | 4660.63 |

**Supplementary table 3.** Model fits for experiment 2, e-greedy model

| | B ($\mu$) | w ($\mu$) | Log-likelihood |
| --- | --- | --- | --- |
| MB | 0.144 | x | -354.3186842 |
| SR | 0.118 | x | -359.19746 |
| MB+SR | 0.056 | 0.481 | -254.18709 |

### Hybrid model: the balance between SR vs. MB parameters

We further analyzed SR-MB weighting parameters to assess the balance between the MB and SR systems using a hierarchical model. Specifically, we estimated the mean and standard deviation of a Gaussian distribution from which individual participant w parameters were drawn, as well as the individual participant parameters. For experiment 1, this produced a group-level w mean of 0.887. Individual participant parameter estimates are displayed in Supplementary figure 4. As noted, the SR effect appears to be smaller in experiment 1 compared to experiment 2. However, the revaluation measures are different in the two studies, study one compares revaluation scores within participants whereas study 2 addresses proportion of participants who changed preference. Therefore, more experiments are required to compare any effect sizes across tasks. That said, in experiment 1 participants were passively exposed to the state transitions rather than exploring them via action selection (as in experiment 2). A consequence of passively exploring the state space was that participants were exposed to states with controlled policies. However, in experiment 2 the agent’s policy was dependent on choices. It is therefore possible that larger signatures of the successor representation would be observed in human behavior in larger state spaces and active selection of actions.

**Supplementary figure 4.**
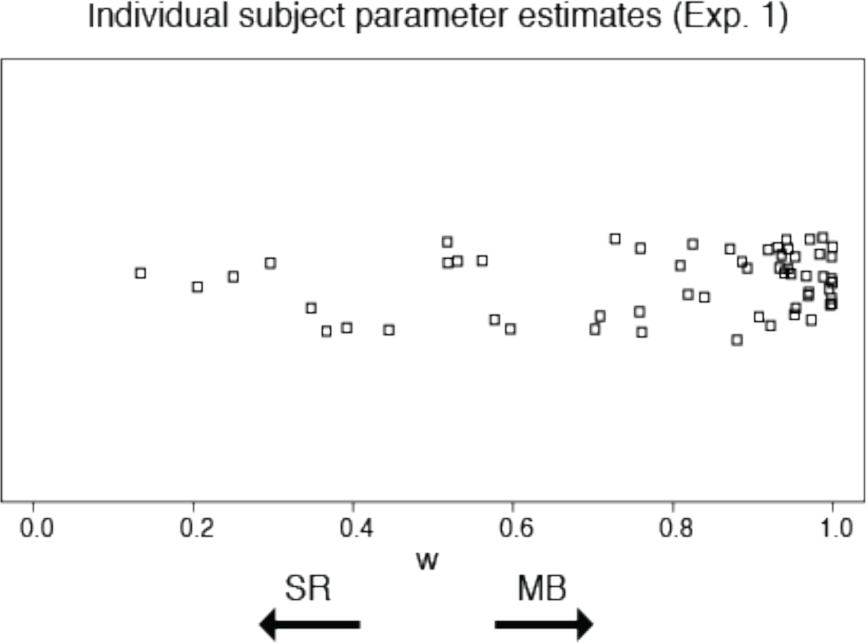
Individual participant parameter estimates from Experiment 1.

### Within-participant consistency of reward and transition revaluation

We investigated within-participant consistency of the transition-reward revaluation asymmetry by computing the effect size (Cohen’s d) for each participant separately. The histogram of effect sizes is shown in Supplementary figure 5.

**Supplementary figure 5.**
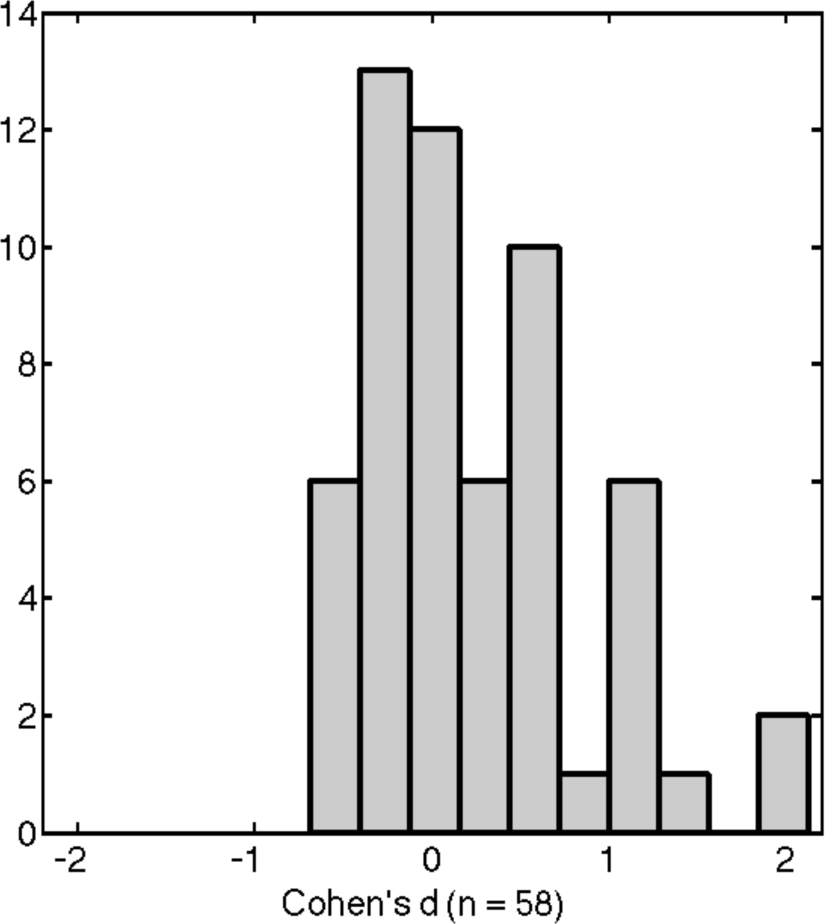
Participant-specific effect size for the transition vs. reward devaluation comparison (paired-sample t-test). Average effect size is 0.24.

### The relationship between learning duration revaluation performance

Supplementary figure 6 plots individual participants’ mean revaluation score against mean length of learning phase. To assess whether this trend is significant, we built a linear model where the dependent variable was mean revaluation score for each participant and the independent variable was the mean length of learning phase for that participant. This revealed a small, yet significant effect of the length of the learning phase on revaluation score (F(1, 56) = 5.05, p = 0.029) with an R-squared of 0.07.

**Supplementary figure 6.**
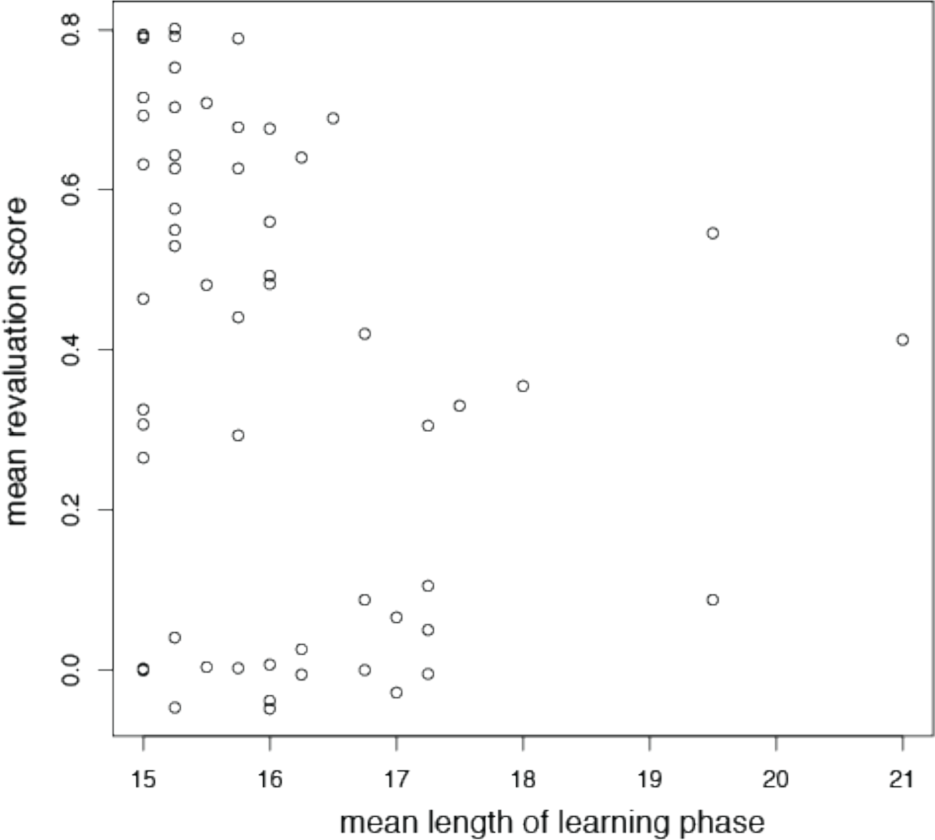
Revaluation score as a function of learning phase length.

Supplementary figure 7 plots individual participants’ mean revaluation score against mean length of revaluation phase. As above, we used a linear model to assess the effect of length of revaluation phase on revaluation score. This analysis revealed a significant effect of length of revaluation phase on revaluation score (F(1, 56) = 4.43, P = 0.04) with R-squared of 0.057.

**Supplementary figure 7.**
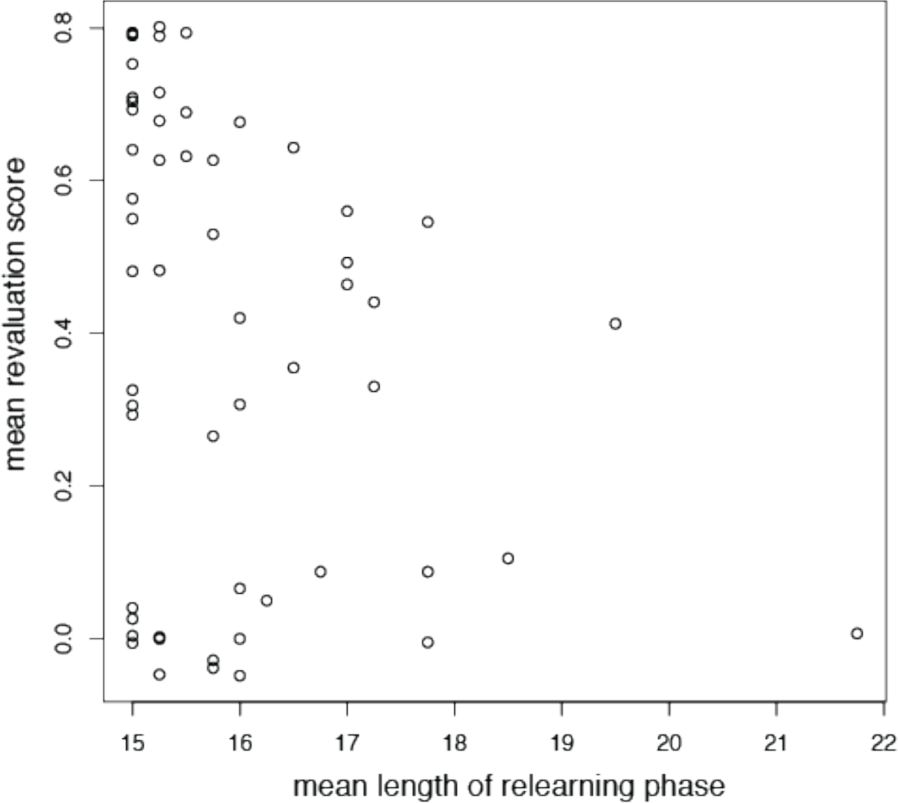
Revaluation score as a function of relearning phase length.

### The relationship between learning duration and reaction time

Supplementary figure 8 plots individual participants’ mean test trial reaction time against their mean length of learning phase. A linear model analysis failed to find a significant effect of length of revaluation phase on test trial reaction time (F(1, 56) = 0.6286, P = 0.4312) with R-squared of -0.006559.

**Supplementary figure 8.**
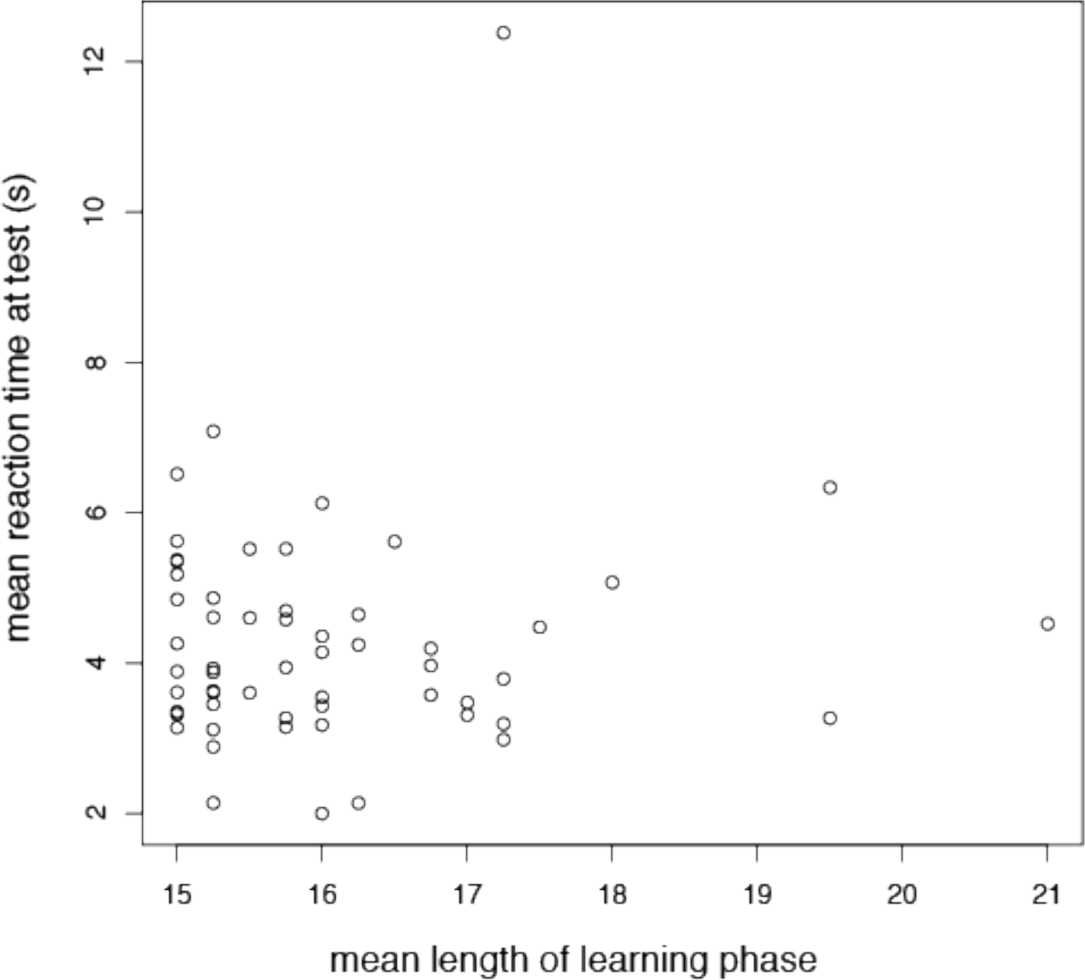
Reaction time during the learning phase.

Supplementary figure 9 plots individual participants’ mean test trial reaction time score against their mean length of relearning phase. A linear model analysis failed to find a significant effect of length of relearning phase on test trial reaction time (F(1, 56) = 0.1437, P = 0.706) with R-squared of -0.01525.

**Supplementary figure 9.**
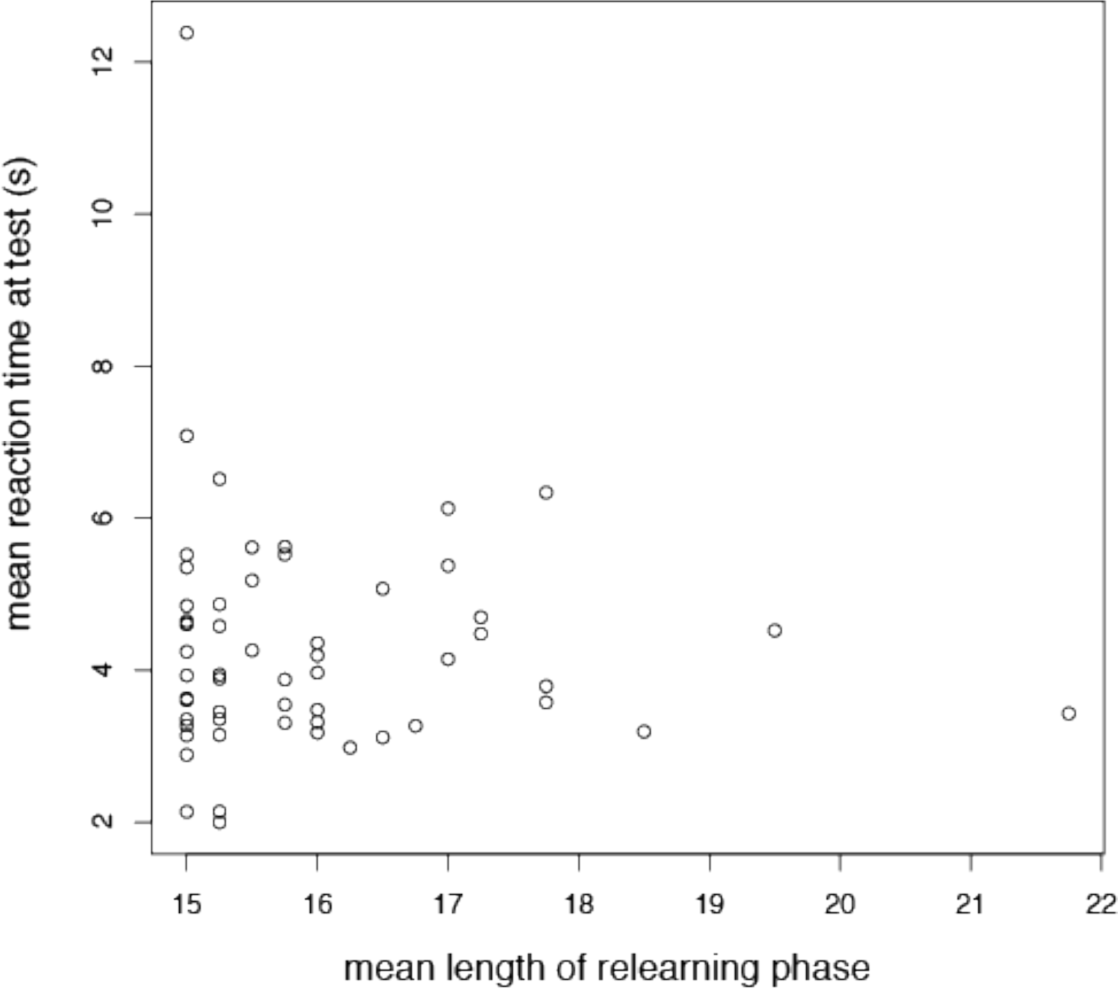
Reaction time during the relearning phase.

## Footnotes

1 Princeton Neuroscience Institute and the Psychology Department, Princeton University

2 Center for Neural Science, NYU

3 Department of Psychological and Brain Sciences, Dartmouth College

4 DeepMind and Gatsby Computational Neuroscience Unit, UCL

5 Department of Psychology and Center for Brain Science, Harvard University

